# Norepinephrine acts through radial astrocytes in the developing optic tectum to enhance threat detection and escape behavior

**DOI:** 10.1101/2025.07.10.664211

**Authors:** Nicholas J. Benfey, Finnley Cookson, David Foubert, Erica Cianfarano, Olivia Ruge, Anton T. Benfey, Anne Schohl, Edward S. Ruthazer

**Affiliations:** Montreal Neurological Institute-Hospital, Department of Neurology and Neurosurgery, McGill University, Montréal, Québec, H3A 2B4, Canada; Information Technology Centre, Department of Computer Science, University of New Brunswick, Fredericton, New Brunswick, E3B 5A3, Canada

## Abstract

The ability to switch behavioral states is essential for animals to adapt and survive. We investigated how norepinephrine (NE) activation of radial astrocytes alters visual processing in the optic tectum (OT) of developing *Xenopus laevis*. NE activates calcium transients in radial astrocytes through α1-adrenergic receptors. NE and radial astrocyte activation shifted OT response selectivity to preferentially respond to looming stimuli, associated with predation threat. NE-mediated astrocytic release of ATP/adenosine reduced excitatory transmission by retinal ganglion cell axons, without affecting inhibitory transmission in the OT. Blockade of adenosine receptors prevented both decreased neurotransmission and the selectivity shift. Chemogenetic activation of tectal radial astrocytes reproduced NE’s effects and enhanced behavioral detection of looming stimuli in freely swimming animals. NE signaling via radial astrocytes enhances network signal-to-noise for detecting threatening stimuli, with important implications for sensory processing and behavior.

## Introduction

Animals navigate complex environments, requiring rapid transitions between cognitive states to optimize context-dependent behavior. Neuromodulators such as serotonin (5-HT), acetylcholine (ACh), dopamine (DA), and norepinephrine (NE) critically alter neural circuit excitability, function, and plasticity, thereby governing brain states and behavior (Marder, 2012; Bargmann and Mader, 2013; Lee and Dan, 2012; Yokogawa et al., 2012; Marques et al., 2020). Glial cells, especially astrocytes, are direct targets of neuromodulators, which serve as crucial mediators of brain state switching (Poskanzer and Yuste, 2016; Mu et al., 2019; Lines et al., 2020). For instance, astrocytic ACh receptors mediate ACh induced plasticity in the visual cortex and hippocampal circadian rhythms in mice (Chen et al., 2012; Navarrete et al., 2012, Papouin et al., 2017). In the mouse amygdala, astrocytic oxytocin receptors shift affective states (Wahis et al., 2021), while in zebrafish and fruit flies, NE or its analogs act through astrocytes to mediate behavioral changes (Ma et al., 2016; Mu et al., 2019).

NE, primarily released by neurons in the locus coeruleus (LC), regulates diverse neural functions including sleep/wake cycles, attention, learning, memory, cortical plasticity, signal-to-noise enhancement, and vigilance/arousal (Bear and Singer, 1986; Berridge and Waterhouse, 2003; Sara, 2009; Aston-Jones and Cohen, 2005). NE is also crucial for stress/emotional regulation (McCall et al., 2015). Dysfunction of NE signaling in the mouse superior colliculus impairs response inhibition and attention (Mathis et al., 2015). Astroglia, particularly in the visual system, are highly responsive to NE, showing large α1R-mediated calcium increases (Giaume et al., 1991; Kulik et al., 1999; Gordon et al., 2005; Bekar et al., 2008; Ding et al., 2013; Paukert et al., 2014; Bazargani and Attwell, 2017; Agarwal et al., 2017; Slezak et al., 2019; Mu et al., 2019; Oe et al., 2020; Ye et al., 2020; Gray et al., 2021; Wahis et al., 2021b). Some work suggests NE’s effects may be predominantly astrocyte-mediated (Mu et al., 2019; Ye et al., 2020). While NE robustly activates visual system astrocytes, its specific effects on visual circuit processing via astrocytes remain unclear.

The optic tectum (OT), or superior colliculus in mammals, integrates sensorimotor information and processes complex visual information (Isa et al., 2021, Li et al., 2022). In fish and frogs, radial astrocytes tile the brain’s surface, resembling radial glial progenitors but also exhibiting mature astrocyte properties thus their recent designation as radial astrocytes (Kriegstein and Alvarez-Buylla, 2009; Tremblay et al., 2009; Sharma and Cline, 2010; Sild et al., 2016; Mu et al., 2019; Benfey et al., 2021). To understand how NE-activated astrocytes shape visual processing during early development, we used *in vivo* GCaMP6s imaging in the *Xenopus laevis* retinotectal system (Chen et al., 2013; Benfey et al., 2021). In this system, retinal ganglion cell (RGC) axons project to the contralateral OT and synapse onto tectal neurons (Figure 1A). Our previous work shows radial astrocytes are dynamic, and crucial for synaptic maturation in the OT (Tremblay et al., 2009; Sild et al., 2016; Van Horn et al., 2017; Benfey et al., 2021). Using a combination of visual stimulation and acute pharmacological manipulation of the OT in albino GCaMP6s expressing tadpoles, together with semi-automated ROI extraction using Suite2p (Pachitariu et al., 2017), we have observed and analyzed neuron-glia communication in the OT *in vivo* (Benfey et al., 2021). Radial astrocyte endfeet are easily identifiable structures (Figure 1B), which facilitates segmentation and quantification of the overall calcium activity occurring in radial astrocytes in the OT (Benfey et al., 2021)

**Figure 1.**
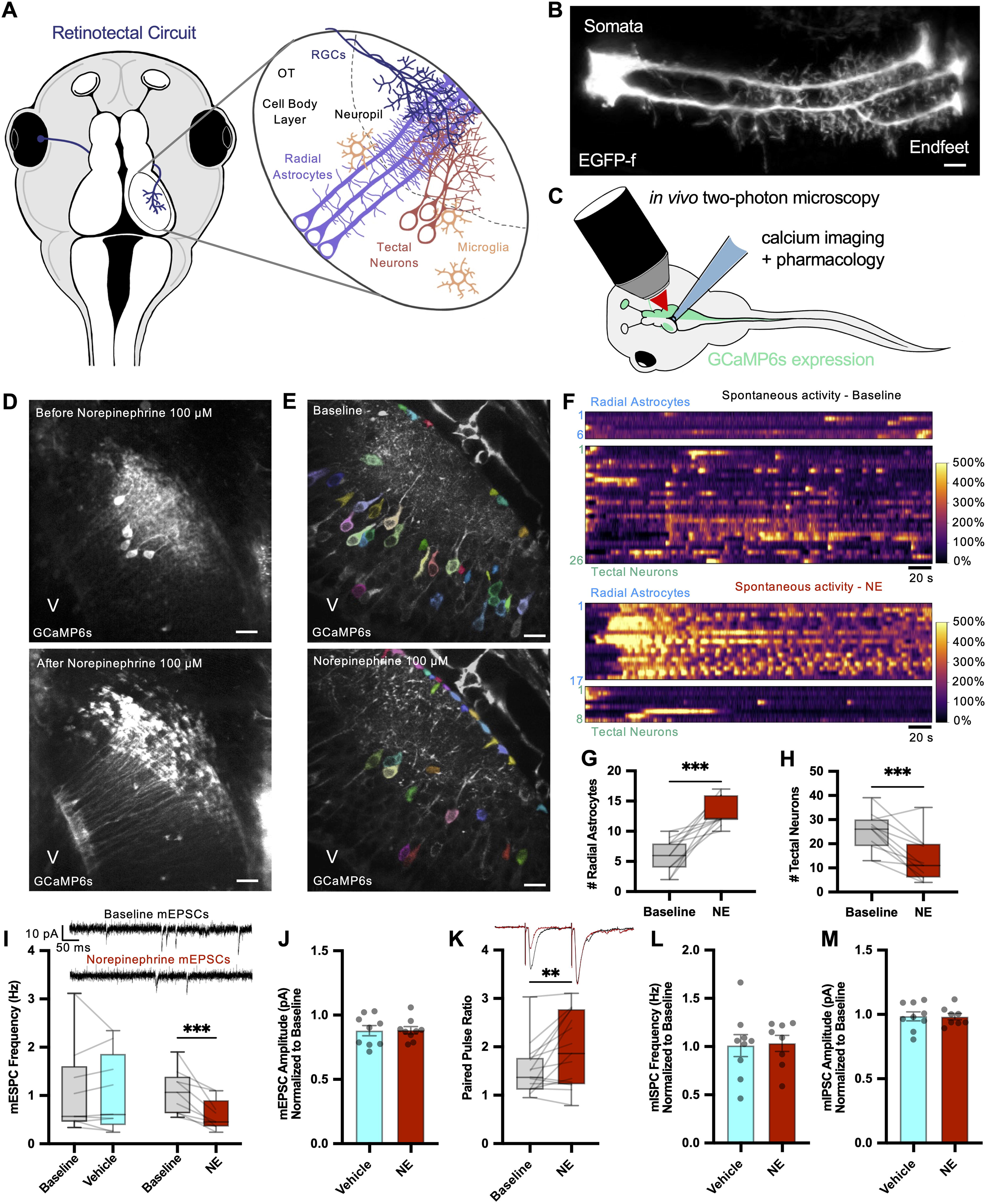
Norepinephrine increases radial astrocyte activity and decreases neuronal activity in the optic tectum. **(A)** *Xenopus laevis* retinotectal projection: RGC axons innervate the contralateral OT. Cellular organization shown. **(B)** Two EGFP-expressing radial astrocytes. Scale: 10 µm. **(C)** Experimental setup schematic. **(D)** GCaMP6s z-projection through 150 µm depth in the right OT hemisphere after TTX (1µM) wash-on (top), and subsequent NE (100 µM) application (bottom). V: tectal ventricle. Scale: 40 µm. **(E)** Average GCaMP6s signal in a single optical section during baseline (top) and after NE (100 µM) (bottom). Color indicates ROIs of tectal neurons and radial astrocyte endfeet idendified by Suite2p. Total time 300s. Scale: 20 µm. **(F)** Baseline normalized (ΔF/F_0_) spontaneous activity traces of radial astrocytes and tectal neurons, baseline (top) and post-NE (100 µM) (bottom). **(G, H)** Number of active radial astrocytes increased (G) and tectal neurons decreased (H) after NE application (single optical section, 5 min). **(I)** Representative whole-cell mEPSC recordings from tectal neurons in a whole brain preparation (top). NE induced a decrease in mEPSC frequency (bottom). **(J)** mEPSC amplitude was unaffected. **(K)** Representative retinotectal EPSCs evoked by paired-pulse stimulation at the optic chiasm (top) NE treatment increased PPR (EPSC2/EPSC1). **(L)** mIPSC frequency and **(M)** amplitude was unaffected by NE treatment. See Table S1 for statistical details.

Here, we show that NE activates tectal radial astrocytes via α1Rs, leading to ATP/Adenosine-dependent reduction in excitatory presynaptic input. NE shifts the OT towards encoding specific, threat-related stimuli like looming cues, which elicit escape behaviors (Barker and Baier, 2015; Temizer et al., 2015; Abbas et al., 2017; Lim and Ruthazer, 2021). Direct chemogenetic activation of tectal radial astrocytes replicated these effects and increased loom detection probability in freely swimming animals, while decreasing exploratory behavior. Our results suggest NE signals through α1Rs on radial astrocytes in the OT to reduce excitatory presynaptic input, and biasing the tectum towards threat detection, underscoring the importance of astrocytes in mediating neuromodulatory effects.

## Results

### Radial astrocytes in the Xenopus laevis optic tectum are directly responsive to norepinephrine

To determine if NE directly activates radial astrocytes, we performed resting state GCaMP6s imaging in the OT before and after NE application (100 µM) with a small incision above the tectal ventricle for facilitated drug access (Figure 1C) (Benfey et al., 2021). Basal calcium activity in tectal neurons and radial astrocyte endfeet was recorded for 5 min before (Figure 1D Top) and after NE addition (Figure 1D Bottom) in the presence of TTX (1µM) to eliminate neuronal action potentials (Video S1). NE dramatically elevated calcium in nearly all radial astrocytes.

### Spontaneous and evoked neural activity in the optic tectum are reduced by norepinephrine

Without TTX, NE (100 µM) significantly increased the number of spontaneously active radial astrocytes (Figure 1E,F,G) and significantly decreased spontaneously active tectal neurons (Figure 1E,F,H)(Video S2). This result confirms reports of an inverse neuron-glia calcium relationship in response to NE in zebrafish (Mu et al., 2019).

Whole-cell patch clamp recordings from tectal neurons showed NE (100 µM), but not vehicle, significantly reduced miniature excitatory postsynaptic current (mEPSC) frequency (Figure 1I) without affecting amplitude (Figure 1J). Reduced mEPSC frequency with no amplitude change suggests reduced presynaptic release probability. Consistent with this, electrical stimulation of RGC axons at the optic chiasm revealed a significant increase in the paired-pulse ratio (PPR) in NE-treated (Figure 1K), but not vehicle-treated (Figure S1A), animals. NE did not alter tectal neuron membrane potential (Figure S1B) or number of spikes evoked by current injection (Figure S1C), indicating no change in intrinsic excitability. Furthermore, NE did not change mIPSC frequency (Figure 1L, S1D,E,F) or amplitude (Figure 1M, S1D,G,H). These data suggest that reduced presynaptic glutamate release from RGC axons may underlie the NE-mediated reduction in OT neuronal activity.

### Norepinephrine selectively alters visual processing and increases the saliency of looming stimuli in the optic tectum

We assessed NE’s effects on visually-driven OT activity by presenting stimuli on an LCD display (Figure 2A) (Benfey et al., 2021, Li et al., 2022). We first presented 25 small, randomly drifting black dots (∼10°, 1s; “dots”) mimicking debris/food (Barker and Baier, 2015; F rster et al., 2020). After 20s, a looming stimulus (”loom”; small black circle expanding to fill screen in 1s), mimicking predation was presented (Lim and Ruthazer, 2021). Dots and loom stimuli were alternated (7 presentations each over 5 min) during baseline and again after NE application (Figure 2B)(Video S3). NE significantly increased the number of active radial astrocytes (Figure 2C) and decreased active neurons (Figure 2D). The mean correlation between the activity patterns of tectal neurons significantly increased with NE (Figure 2E), suggesting a more tightly tuned population.

**Figure 2.**
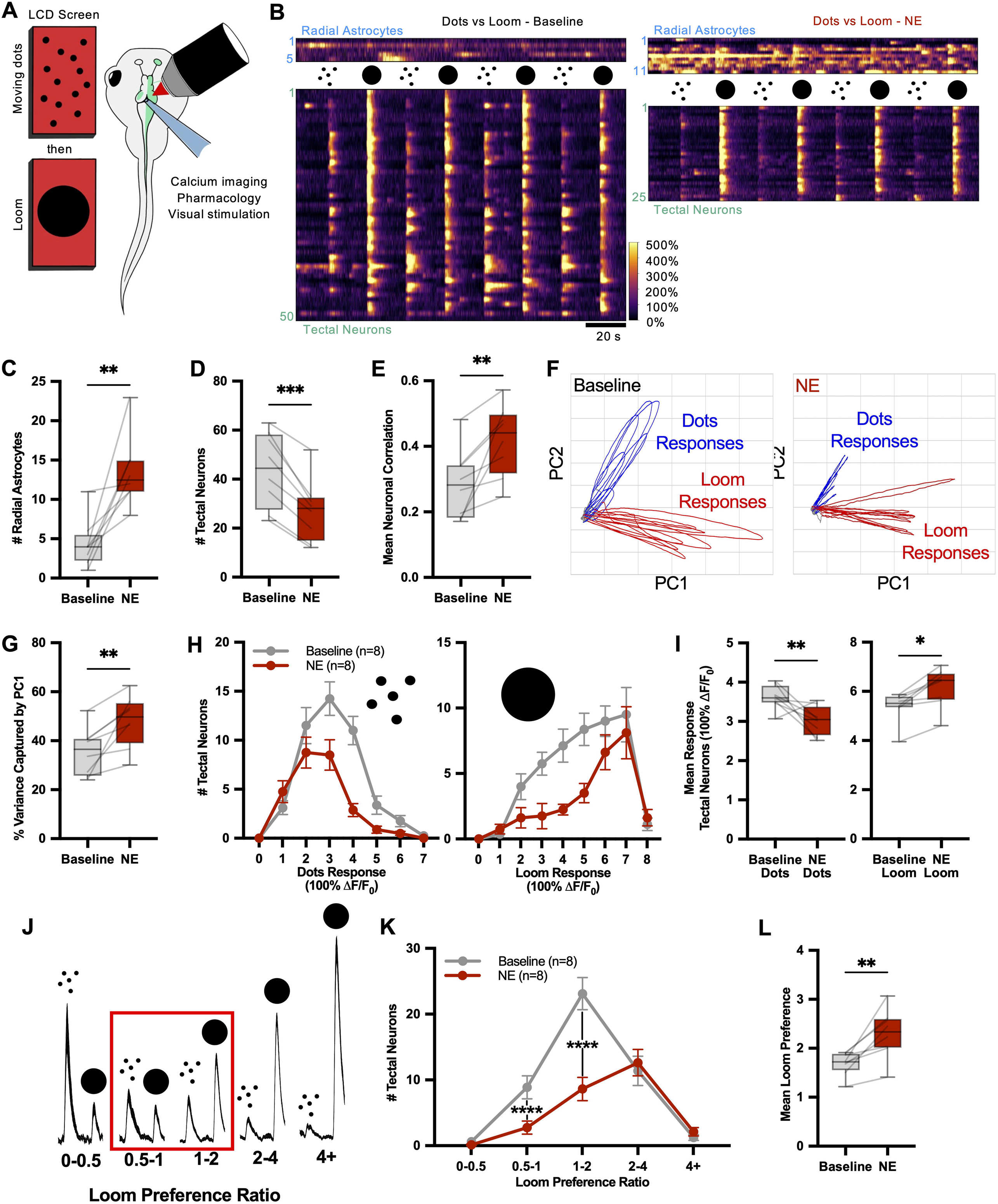
Norepinephrine leads to selective visual processing in the optic tectum. **(A)** Experimental setup schematic. **(B)** Baseline normalized (ΔF/F_0_) calcium activity traces (top: radial astrocytes, bottom: tectal neurons) at baseline (left) and post-NE (100 µM) (right). **(C, D)** Number of active radial astrocytes increased (C) and tectal neurons decreased (D) with NE application. **(E)** Pearson correlation across active tectal neurons increased in NE. **(F)** PCA plots of population activity captured by PC1 and PC2 during baseline (left) and NE (right). Blue: moving dots; Red: looming stimuli. **(G)** A greater percent of total variance was captured by PC1 after NE application. **(H)** Histogram of average peak response amplitudes to dots (left) and loom (right) in tectal neurons. **(I)** Mean peak response amplitudes decreased for dots (left) and increased for loom (right). **(J)** A Loom Preference Ratio (LPR) was calculated per cell as the ratio of response to loom over dots. **(K)** Tectal neuron LPR was shifted in favor of loom stimuli by NE, **(L)** resulting in an increased mean LPR per animal. See Table S2 for statistical details

Principal component analysis (PCA) of population activity (Figure 2F) (Pang et al., 2016; Stringer et al., 2019a; Stringer et al., 2019b) showed PC1 predominantly represented visual stimulus-driven variance, while PC2 captured stimulus type (dots vs. loom). NE significantly increased the percent of variance captured by PC1 (Figure 2G), suggesting NE yields a less complex, more correlated visual representation, potentially facilitating threat detection (Stringer et al., 2019a). Analysis of mean peak neuronal responses (Figure 2H) showed NE decreased high-amplitude responses to dots (Figure 2I, left) but not low-amplitude responses, reducing overall mean dot response. Conversely, NE decreased low-amplitude loom responses but not high-amplitude ones, increasing overall mean loom response (Figure 2I, right). This asymmetry suggests NE increases loom representation by filtering out dot representation. A ‘loom preference ratio’ (LPR; mean peak loom response / mean peak dots response) quantified this shift (Figure 2J). NE treatment caused a dramatic shift to higher LPRs, reducing neurons with dot or weak loom preference (Figure 2K). Mean OT LPR significantly increased with NE (Figure 2L). These results show NE induces an asymmetric, selective alteration of OT visual processing correlated with radial astrocyte activation, leading to a sparse, robust representation of threat-related information.

### Preference for small moving dots with coherent motion in the norepinephrine state

To control for differences in spatial statistics between dot and loom stimuli, we presented alternating dot stimuli (2s duration) with equivalent features but contrasting motion coherence: no coherence (Dots C0) or full coherence (Dots C1) (Figure S2A). NE significantly increased radial astrocyte activity (Figure S2B) and decreased tectal neuron activity (Figure S2C). Unlike dots vs. loom, mean neuronal correlation (Figure S2D) and PC1 variance (Figure S2E) did not increase with NE, suggesting Dots C1 and loom are encoded differently. Mean responses trended towards decreased Dots C0 response and increased Dots C1 response, but neither was significant (Figure S2F,G). NE significantly decreased neurons preferring Dots C0 and Dots C1 (Figure S2H,I), but the reduction for Dots C0 preference was twice that for Dots C1, despite similar baseline numbers. This resulted in a significant increase in mean OT preference for Dots C1 with NE (Figure S2J).

### Alpha-1 adrenergic receptor activation is required for norepinephrine-induced changes in visual response properties of the optic tectum

α1Rs underlie NE-induced calcium elevations in mammalian astrocytes (See comprehensive review by Wahis and Holt, 2021). To determine if α1Rs mediate NE-induced changes in radial astrocytes, we applied NE after pretreating with the α1R antagonist prazosin (PRZ, 50 µM) during the dots vs. loom paradigm (Figure 3A). In PRZ-pretreated animals, NE did not increase radial astrocyte activity (Figure 3B) nor decrease tectal neuron activity (Figure 3C), suggesting both effects require α1R activation. PRZ-treated animals showed higher baseline mean neuronal correlation (Figure 3D), indicating less prominent dot vs. loom selectivity, as predicted if NE acts via α1Rs to enhance loom representation. PC1 variance did not increase in PRZ+NE animals (Figure 3E). The PRZ group showed no NE-induced shifts in dot/loom response preference (Figure 3F) or LPR (Figure 3G,H). These results strongly suggest NE activates radial astrocytes via α1Rs.

**Figure 3.**
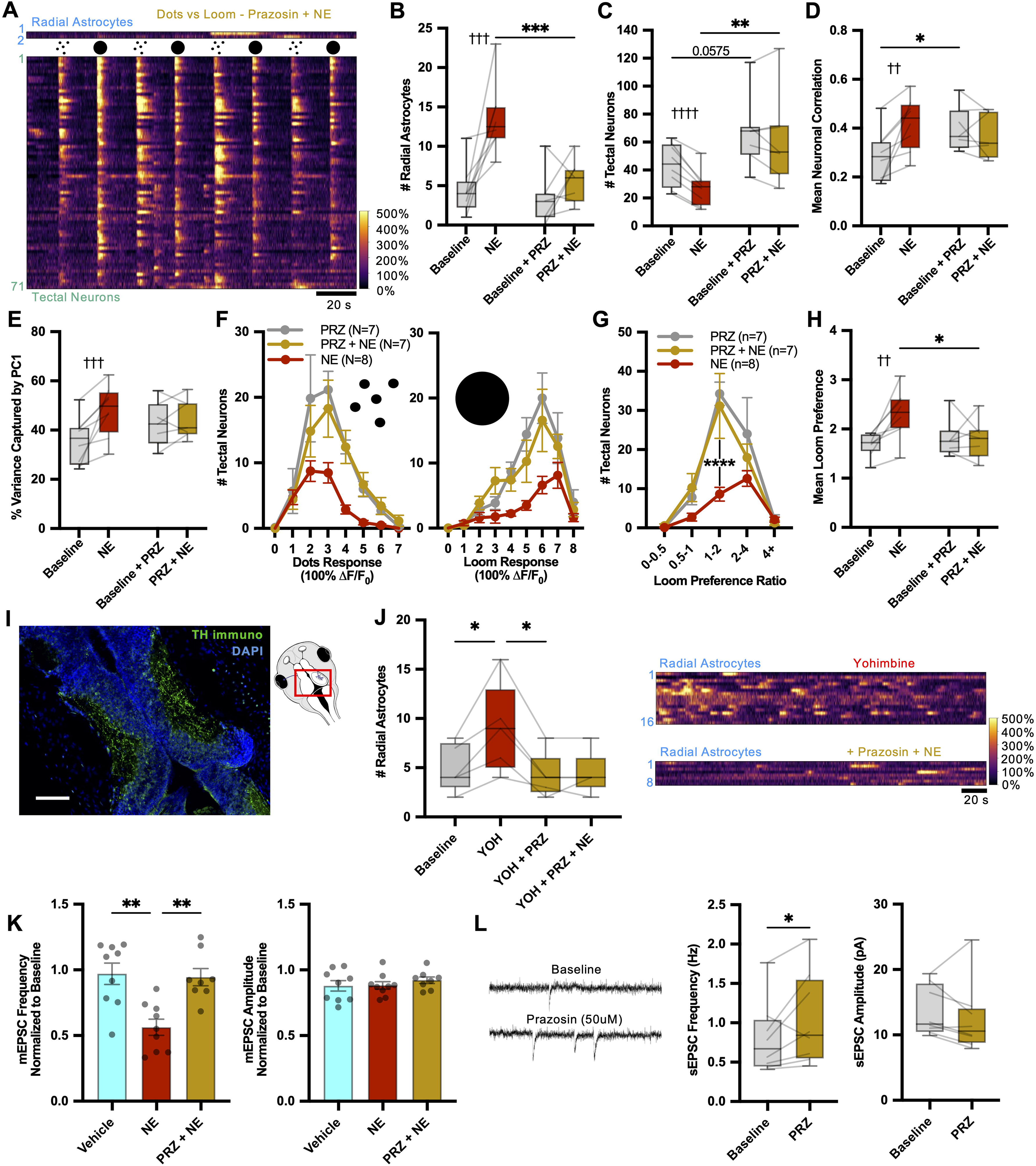
Alpha-1 adrenergic receptor activation is required for norepinephrine-induced changes in selective visual processing. (**A**) Baseline normalized (ΔF/F_0_) calcium traces (top: radial astrocytes; bottom: tectal neurons) during alternating dots/loom stimulation after adding NE (100 µM) under α1-adrenergic receptor inhibition by prazosin (PRZ, 50 µM). **(B, C)** PRZ prevents the increase in active radial astrocytes (B) and the decrease in active tectal neurons (C). **(D, E)** Increase in tectal neuron correlation (D) and population response variance encoded in PC1(E) are also blocked by PRZ. **(F)** More neurons responded both to dots (left) and to loom (right) under PRZ. (NE: n=8 animals, PRZ(+NE): n=7 animals), but (**G, H**) the NE-mediated shift in LPR favoring loom stimuli was prevented in PRZ. **(I)** TH (*green*) immunostaining with DAPI counterstain (*blue*) of OT cryosection. Scale bar: 100 µm. **(J)** α2R antagonist yohimbine (YOH, 20 µM) to elevate endogenous NE release increased the number of active radial astrocytes (top). This was reversed by PRZ, which further blocked the effects of exogenous NE (*left*). Representative radial astrocyte calcium traces (*right*) in YOH, before (*top*) and after (*bottom*) PRZ and NE. **(K)** NE-mediated decrease in mEPSC frequency was blocked by PRZ treatment, with no effect on amplitudes **(L)** Whole-cell recordings of spontaneous synaptic events (without TTX) revealed increased frequency (but not amplitude) after adding PRZ to prevent endogenous α1R activation of astrocytes. See Table S3 for details on statistical tests.

Tyrosine hydroxylase (TH), the enzyme required for synthesis of catecholamines like NE, is expressed in LC noradrenergic projection neurons. TH immunohistochemistry confirmed extensive TH-positive axon ramification in stage 48 tadpole OT neuropil (Figure 3I). Presynaptic α2Rs on noradrenergic terminals are believed to negatively regulate NE release (Paton and Visi, 1969). Applying the α2R antagonist yohimbine (YOH, 20 µM) to elevate endogenous NE release significantly increased radial astrocyte calcium events, similar to exogenous NE (Figure 3J) (Goldberg et al., 1983). α1R blockade with PRZ eliminated YOH’s effects, as well as that of subsequent exogenous NE application. NE-induced mEPSC frequency reduction was also blocked by PRZ (Figure 3K). Bath application of PRZ (without TTX) significantly increased spontaneous EPSC (sEPSC) frequency but not amplitude (Figure 3L), consistent with ongoing endogenous α1R activation in the OT. While α2R blockade might also increase serotonin (5-HT) release (Scheibner et al., 2001), and elicit 5-HT mediated effects on visual processing (Yokogawa et al., 2012; Udoh et al., 2024) acute 5-HT application had no effect on tectal responses to dots/loom stimuli (Figure S3).

### Norepinephrine-induced changes in visual stimulus selectivity require ATP/adenosine signaling

Astrocytic ATP release regulates diverse neural circuits and behaviors (Pascual et al., 2005; Gourine et al., 2010; Boddum et al., 2016; Fujii et al., 2017; Tan et al., 2017; Covelo and Araque, 2018; Eersapah et al., 2019; Agostinho et al., 2020; Corkrum et al., 2020). Its breakdown product, adenosine (ADO), also regulates astrocyte-mediated effects via presynaptic A1, A2a, and A2b receptors (Figure 4A), often by reducing vesicular release (Lopes et al., 2002; Moore et al., 2003; Nasrallah et al., 2024; Shindou et al., 2002). In Xenopus OT, ADO application significantly reduced mEPSC frequency (Figure 4B,C) and increased retinotectal PPR (Figure 4D,E), indicating reduced RGC presynaptic release probability. Pretreatment with AMPCP (5’-nucleotidase inhibitor preventing ATP to ADO conversion) blocked NE’s effect on mEPSC frequency (Figure 4F). Among A1 (DPCPX), A2a (SCH 58261), or A2b (MRS 1754) receptor inhibitors, only the A2a antagonist SCH 58261 prevented NE-induced mEPSC frequency reduction (Figure 4F); amplitudes were unaffected (Figure 4G). This indicates NE-activated astrocytes release ATP, broken down to ADO, which reduces synaptic release via A2a receptors.

**Figure 4.**
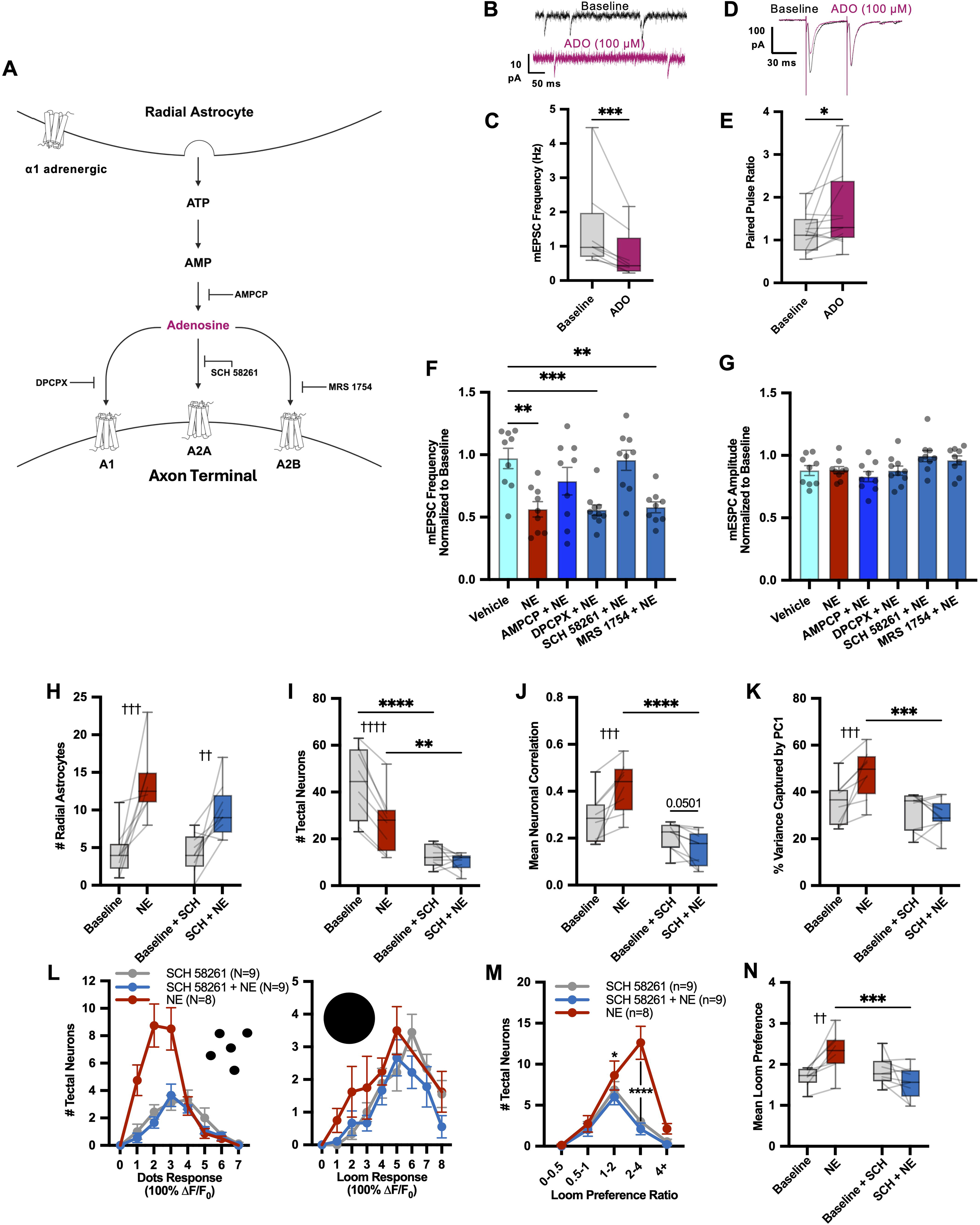
Norepinephrine-induced visual stimulus selectivity changes require ATP/adenosine signaling. **(A)** ATP/adenosine signaling downstream of astrocyte activation. **(B)** Example mEPSC traces before and after application of ADO (100 µM). **(C)** mEPSC frequency is reduced by ADO treatment. **(D)** Representative retinotectal evoked PPR traces with ADO addition. **(E)** ADO increases PPR (EPSC2/EPSC1). **(F, G)** mEPSC frequencies (F) and amplitudes (G) in vehicle, NE (100 µM), AMPCP (100 µM), DPCPX (100 nM), SCH 58261 (100 nM), MRS 1754 (100 nM). Inhibition of AMP degradation to ADO and blockade of adenosine A2a receptors (but not A1 or A2b receptors) prevent the decrease in mEPSC frequency by NE. **(H)** In the presence of SCH 58261, NE still can increase the number of active radial astrocytes, but (**I**) the NE-mediated decrease in active tectal neurons is prevented. **(J)** The NE-mediated increase in correlation between active tectal neurons and (**K**) the increase in variance represented by PC1 are blocked by SCH 58261. **(L)** SCH 58261 inhibits the NE-dependent shift in response amplitudes to dots (*left*) and loom (*right*). (NE: n=8 animals; SCH 58261(+NE): n=9 animals). **(M,N)** The NE-mediated shift in LPR favoring loom is completely prevented in SCH 58261. See Table S4 for statistical details.

To test if NE-driven visual preference changes also depend on A2a signaling, we imaged OT calcium responses to dots/loom with NE after SCH 58261 pretreatment. NE still activated radial astrocyte calcium transients (Figure 4H), but neuronal activity reduction was absent (Figure 4I). A2aR inhibition prevented NE-induced elevated neuronal response correlation and dimensionality reduction (Figure 4J,K). Importantly, NE-mediated increases in loom selectivity were completely prevented by SCH 58261 (Figure 4L-N), despite significant astrocyte activation.

### Blockade of adenosine type 1 receptors or hemichannels also modulates the tectal response to norepinephrine

Astrocytic ATP/adenosine modulatory mechanisms are complex (Bazargani and Attwell, 2016), with A1Rs (Florian et al., 2011; Schmitt et al., 2012; Cao et al., 2013; Yang et al., 2015; Corkrum et al., 2020; Li et al., 2020; Roberts et al., 2022) and connexin-containing gap junctions (GJs)/hemichannels (HCs) also implicated (Lalo et al., 2014; Bazargani and Attwell, 2016; Agostinho et al., 2020; Bennett et al., 2003; Orellana and Stehberg, 2014; Roux et al., 2015; Meunier et al., 2017; Fukuyama et al., 2020; Wahis et al., 2021a; Ribot et al., 2021). To investigate further, we blocked A1Rs (DPCPX, 100 nM) or GJs/HCs (carbenoxolone CBX, 100 µM) before NE application during visual stimulation (Figure S4A). DPCPX or CBX pretreatment did not prevent NE from significantly increasing radial astrocyte activity (Figure S4B) and decreasing tectal neuron activity (Figure S4C). However, unlike NE alone, mean neuronal correlation (Figure S4D) and PC1 variance (Figure S4E) did not significantly increase. This suggests that while visual representation size shrinks, its complexity doesn’t decrease as with NE alone, possibly due to persistent dot responsiveness (Figure S4F). Despite reductions in weakly responsive neurons, the effect was no longer biased to favor loom over dots. Thus, NE did not increase LPR in DPCPX or CBX-treated animals (Figure S4G-I). These results indicate A1Rs and HCs also contribute to NE-induced tectal processing shifts.

### Direct chemogenetic activation of radial astrocytes in the optic tectum is sufficient to mimic norepinephrine-like changes in neuronal activity

In mammals, the TRPV1 receptor is a non-specific cation channel that detects increases in temperature and is potently activated by capsaicin (mTRPV1, EC50: ∼ 3 μM) (Ohkita et al., 2012; Chen et al., 2016). However, endogenous amphibian TRPV1 channels are largely insensitive to capsaicin (Xenopus laevis TRPV1, EC50: ∼ 85.4 μM). Consequently, mTRPV1 expressed in radial astrocytes can be activated with low doses of capsaicin that have no effect on endogenous TRPV1 activity in these animals (Mu et al., 2019). We utilized this approach to target the radial astrocytes of the *X. laevis* OT through targeted electroporation of mTRPV1red plasmid injected into the ventricular space between the hemispheres of the OT, which results in the sparse transfection of the radial astrocytes, whose somata line the periventricular surface (∼10-20 radial astrocytes transfected in each hemisphere of the OT), without transfecting neurons, which are located further from the ventricle in the developing X. laevis OT (Figure 1A, 5A,B) (Tremblay et al., 2009; Sild et al., 2016). Transfected radial astrocytes in the OT can then be selectively activated through application of capsaicin.

To directly test the astrocytes contribution to state switching in the OT, we expressed capsaicin-sensitive mTRPV1(E600K)-tagRFP (mTRPV1red) in radial astrocytes in the OT via electroporation (Figure 1A, 5A,B), allowing selective activation with low dose capsaicin. Whole-cell electrophysiological recording from tectal neurons in a whole brain *ex vivo* preparation from these animals revealed a significant reduction in tectal neuron mEPSC frequency in response to capsaicin wash-on (Figure 5C), comparable to NE’s effect (Figure 1I); mEPSC amplitude was unchanged (Figure 5D).

**Figure 5.**
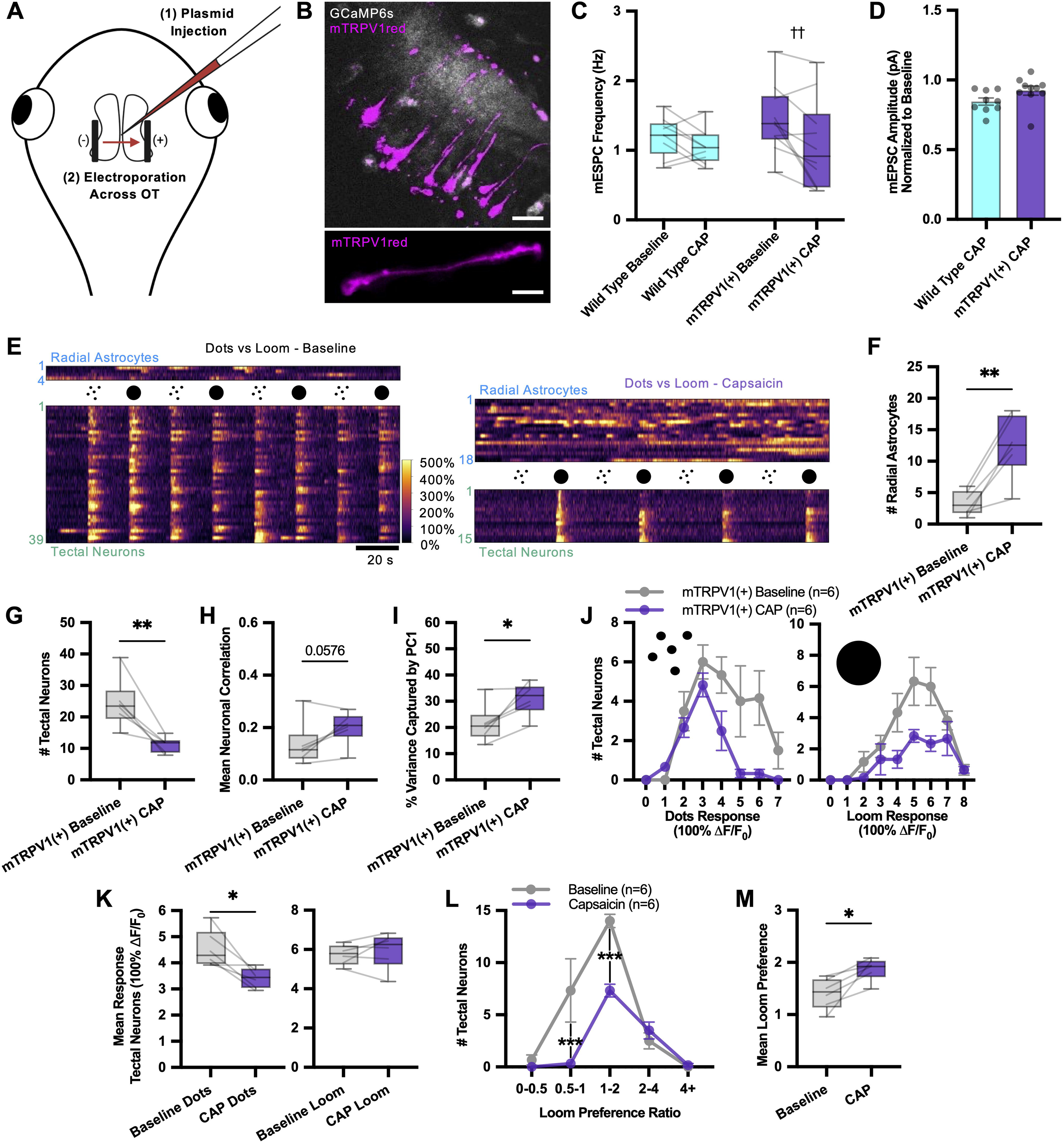
Chemogenetic radial astrocyte activation mimics norepinephrine-like neuronal activity changes and enhances looming stimuli detection. **(A)** Electroporation of mTRPV1red into OT radial astrocytes. **(B)** Top: 2P optical section showing mTRPV1red-transfected radial astrocytes (*magenta*) in GCaMP6s-expressing (*gray*) OT. V: tectal ventricle. Bottom: z-projection of single mTRPV1red-transfected radial astrocyte. Scale: 20 µm. **(C)** Capsaicin (CAP 10 µM) application reduces mEPSC frequency in whole cell recordings from tectal neurons in mTRPV1red-transfected, but not in untransfected, animals. (**D**) Amplitudes are unchanged by CAP wash-on. **(E)** OT radial astrocyte and neuronal calcium traces during dots/loom stimulation (baseline, *right*; post-capsaicin, *left*). **(F)** CAP activates radial astrocytes and (**G**) reduces the number of responsive tectal neurons. **(H)** Pearson correlation between active tectal neurons shows a positive trend with CAP application and (**I**) increased percent variance represented by PC1. **(J)** Average peak response amplitudes to dots (left) and loom (right) reveal a loss of strong dot responses, resulting in (**K**) a reduction in peak response amplitudes for dots (left) but not loom (right). **(L)** CAP treatment results in a rightward LPR shift in neuronal responses, favoring loom stimuli, and (**M**) increased mean LPR per animal. See Table S5 for statistical details.

In GCaMP-expressing animals transfected with mTRPV1red in radial astrocytes, capsaicin (10 µM) application (Figure 5E) activated both mTRPV1red-expressing astrocytes and untransfected neighbors, suggesting propagated activation via diffusible signals or gap junctions. Selective astrocyte activation significantly increased active glia (Figure 5F) and decreased active tectal neurons (Figure 5G). Neuronal correlation increased in 5/6 animals (Figure 5H), and PC1 variance significantly increased (Figure 5I), suggesting astrocyte activation shifts the OT to a lower-dimensional, robust encoding state similar to NE (Figure 2). Chemogenetic astrocyte activation decreased strong, but not weak, dot responses (Figure 5J,K), similar to NE (Figure 2H,I). Consequently, the number of neurons with low LPRs (dot preference) decreased, while strong loom preference neurons were unaffected (Figure 5L), significantly increasing mean OT LPR (Figure 5M). These results strongly suggest radial astrocytes mediate NE-induced visual information filtering.

### Chemogenetic activation of radial astrocytes in the optic tectum reduces exploratory behavior and increases escape behavior in freely swimming animals

Our findings suggest improved loom detection as a result of radial astrocyte activation. We assessed behavioral effects of radial astrocyte activation in the OT by presenting loom stimuli (10 times, 20s ISI) to freely swimming mTRPV1red-expressing animals, tracking exploratory and escape behaviors (Figure 6A)(Video S4,5). Capsaicin-treated untransfected animals and vehicle-treated mTRPV1red animals showed similar loom-evoked escape rates and exploratory behavior levels (Figure 6B,C). Capsaicin-induced chemogenetic activation of radial astrocytes in mTRPV1red animals significantly decreased exploratory behavior (Figure 6B) and increased loom-evoked escape rates (Figure 6C). This suggests astrocyte activation induces heightened vigilance, increasing freezing and escape probability, likely by enhancing loom feature prominence through suppressed weak/spontaneous neuronal OT activity. While not excluding additional NE effects on neurons, our chemogenetic experiments indicate NE’s predominant OT effects are astrocyte-mediated.

**Figure 6.**
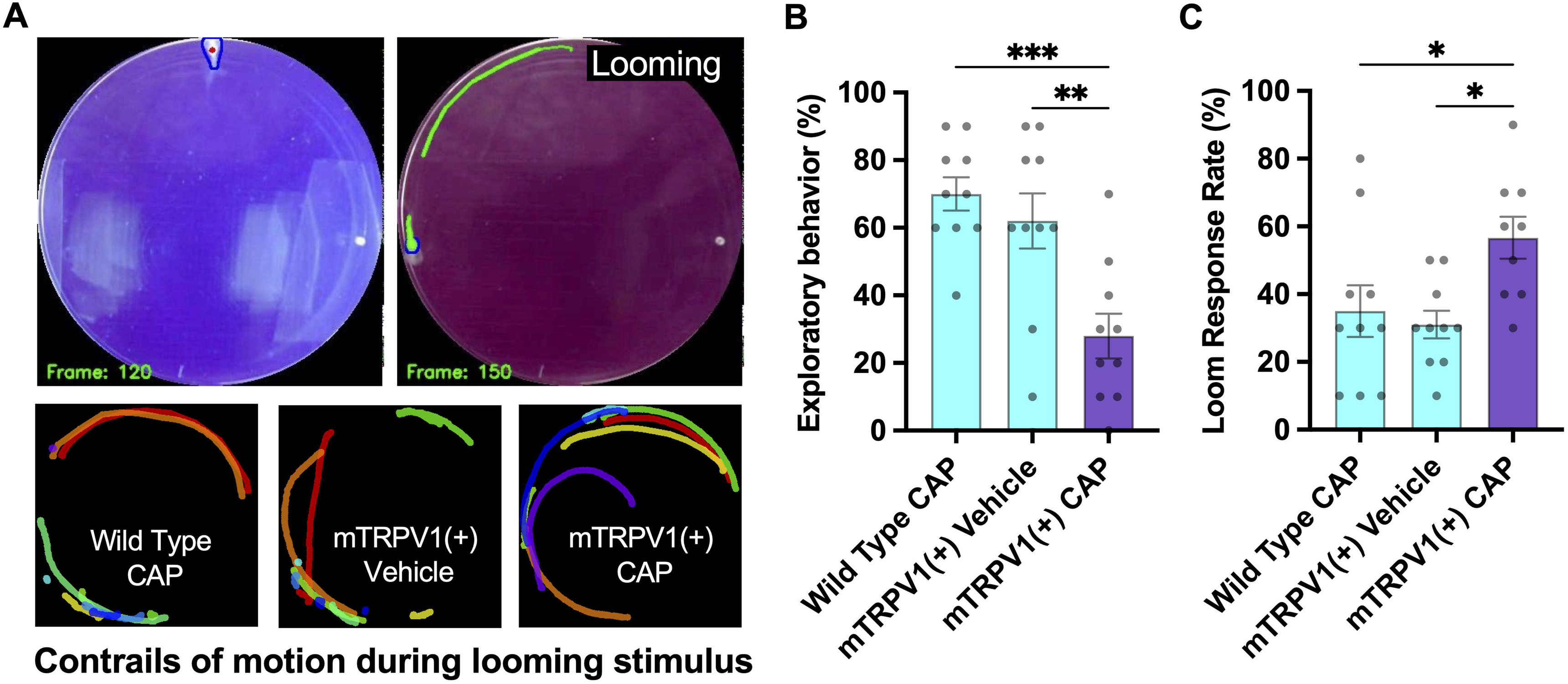
Chemogenetic radial astrocyte activation reduces exploratory behavior and increases escape behavior in freely swimming animals. **(A)** Free-swimming tadpole in a Petri dish next to a screen onto which visual stimuli were projected for the loom escape assay (before loom, *top left*, during loom, *top right*). Contrails show escape trajectories. Representative escape contrails (10 looms). *Bottom left*: untransfected tadpole + capsaicin (10 µM). *Bottom middle*: mTRPV1red-transfected tadpoles + vehicle. *Bottom right*: mTRPV1red + capsaicin (10 µM). **(B)** Exploratory behavior was quantified by measuring the fraction of time the tadpole spent actively locomoting during the inter-stimulus period. (**C**) Loom-evoked escape behavior was scored by counting the percent of loom presentations (10 per animal) that evoked an escape response. Chemogenetic activation of radial astrocytes resulted in a large increase in escape probability, despite reduced overall exploratory locomotion. See Table S6 for statistical details.

## Discussion

This study revealed a fundamental role for radial astrocytes in NE-mediated neuromodulation of brain states. Radial astrocyte activation via α1-adrenergic receptors and subsequent ATP/adenosine release reduces presynaptic release probability, preferentially at excitatory synapses. The number of visually responsive tectal neurons and their stimulus selectivity shifted dramatically to favor salient, threatening stimuli like loom and coherent motion. Direct chemogenetic astrocyte activation largely mimicked NE’s effects, suggesting their essential role. Loom escape experiments show astrocyte-mediated brain state changes directly affect sensorimotor behaviors, suggesting gliotransmission alters perceptual salience.

Our finding that noradrenergic signaling in the optic tectum primarily regulates synaptic transmission through the release of ATP/adenosine from the radial astrocytes activated via α1-adrenergic receptors is consistent with reports in various other models. In the hindbrain of the zebrafish, α1 activation of radial astrocytes has been shown to elevate extracellular ADO levels, which in turn inhibits behavioral passivity (Chen et al., 2025). Curiously, in this hindbrain circuit the inhibition was prevented by pharmacological blockade of A2bRs, but not by blocking A1R or A2aR. Furthermore, similar to our findings at the Xenopus retinotectal synapse, spontaneous and evoked release in the mouse hippocampus are inhibited by ATP/adenosine released in response to α1-adrenergic activation of astrocytes (Lefton et al., 2025). In this case, pharmacological blockade of A1Rs prevents the effect. Thus, while purinergic signaling via astrocyte activation appears to be a highly conserved mechanism, the ADO receptors targeted varies between different brain areas and behavioral outcomes.

In aquatic vertebrates like *Xenopus laevis*, the OT plays a key role in detection of looming stimuli and in driving escape behavior (Temizer et al., 2015; Dunn et al., 2016; Lim and Ruthazer, 2021). The OT mediates approach/avoidance based on stimulus size relative to the eye; small objects trigger approach, larger ones avoidance (Barker and Baier, 2015). Consistent with our findings, critical image size processing is reportedly distributed across tectal neurons, not localized to specific “loom-detectors” (Dunn et al., 2016). Modeling the neural networks processing visual information in the Xenopus laevis OT has also implicated levels of recurrent network activity in loom detection (Jang et al., 2016). Curiously, a group of neurons in the superficial layers of the OT, which exhibit high levels of gap junctional coupling, have been found to control recurrent network activity in the Xenopus laevis OT (Liu et al., 2016).

Additionally, it has been suggested that these neurons may boost the reliability of signals in the deep layers of the OT which are known to be involved with mediating escape behavior. This is particularly relevant to our observations as in both the norepinephrine (NE) state, as well as the state induced by chemogenetic activation of radial astrocytes in the OT, we consistently observed neurons in the superficial and neuropil layers of the OT that remained highly responsive, even as activity in periventricular neurons generally decreased.

Circuit models of tectal neurons have also implicated a population of superficial interneurons (SINs), primarily found embedded within the synaptic neuropil region of the OT, as being important for the processing and filtering of size related visual information transmitted between RGCs and the tectal neurons in the deeper layers of the OT (Del Bene et al., 2010; Barker and Baier, 2013; Temizer et al., 2015; Dunn et al., 2016; Abbas et al., 2017). These SINs extend widely branching dendrites throughout much of the tectal neuropil and contribute to the computations that drive the initiation of escape behaviors (Barker and Baier, 2013; Temizer et al., 2015; Dunn et al., 2016; Barker et al., 2021). It has been shown that SINs enhance the detection of visual stimuli, likely by suppressing smaller distractions in the environment, or by enhancing the ability to distinguish between objects of different sizes in the environment, thereby facilitating the initiation of specific sensory-motor behaviors such as loom-evoked escapes (Barker and Baier, 2015). In both Xenopus laevis and zebrafish, the vast majority of SINs (∼80%) are GABAergic (Miraucourt et al., 2012; Barker et al., 2021; Mu et al., 2019). SINs are highly responsive to loom and often remained active in the NE state, or following chemogenetic activation of radial astrocytes in the OT, when many tectal neurons in the cell body layer of the OT become quiescent. In this context, it is significant that the effects of NE on reducing vesicular release probability that we observed were limited to excitatory synapses and did not impact GABAergic inputs. Occasionally, we observed persistent calcium elevations in these neurons lasting tens of seconds following either the addition of NE or chemogenetic activation of radial astrocytes which strongly suggests that these neurons are likely to play a role in mediating the effects of radial astrocytes on visual processing in the OT as well as the enhanced loom detection that we observe in freely swimming animals. In this context, it is interesting that the effects of NE on reducing vesicular release probability that we observed were limited to excitatory synapses and did not impact GABAergic inputs. Optogenetic activation of SINs in zebrafish OT was sufficient to mediate the shift in swimming behavior observed following the activation of radial astrocytes (Mu et al., 2019). Further experiments aimed at dissecting the circuit mechanisms mediating NE-driven state switching in the OT will require careful investigation of the relationship between radial astrocytes and SIN function and their roles in sensory processing.

Our findings also implicate gap junction/hemichannel opening in the NE-driven, astrocyte-mediated perceptual filtering of dots. NE regulates gap junction formation and connexin 43 redistribution via α1R activation (Giaume et al., 1991; Nuriya et al., 2018). Both GJs and HCs have important astrocytic functions (Orellana and Stehberg, 2014). In the olfactory bulb, astrocytic connexin 43 hemichannels open with glial calcium increases, releasing ATP/adenosine and modulating slow waves (Orellana and Stehberg, 2014; Roux et al., 2015). Calcium-dependent connexin 43 hemichannel opening in astrocytes is also linked to D-serine release, an important gliotransmitter in Xenopus OT (Meunier et al., 2017; Van Horn et al., 2017). In the amygdala, astrocytic connexin 30/43 hemichannels release D-serine, shifting affective states post-oxytocin release (Wahis et al., 2021a). CBX has been reported to prevent gliotransmitter release through connexin 43 hemichannels (Fukuyama et al., 2020). Investigating specific connexins in OT radial astrocytes will be important for understanding their function.

Additionally, tectal neurons silenced by NE/glial activation were often in less mature posterior OT regions, weakly responsive to both dots and loom. Enhanced glutamate clearance by activated radial astrocytes might also contribute to silencing tectal neurons, particularly in less mature regions. Local astrocyte membrane voltage changes alter glutamate clearance efficiency (Armbruster et al., 2022), and α1R activation enhances astrocytic glutamate clearance (Hansson and Ro nnba ck, 1989; Fahrig, 1993). This might silence immature neurons with low signal-to-noise ratios during high vigilance.

Recent studies highlight radial astrocytes’ critical role in NE-mediated behavioral state transitions and OT neuronal activity modulation (Uribe-Arias et al., 2023). Synchronized calcium elevations in zebrafish OT radial astrocytes are LC NE-mediated and modulate tectal neuron direction selectivity and long-distance functional correlations post-escape. These neuron-glia interactions may help the OT adapt to new contextual information, informing subsequent behavioral decisions. This synchronized astrocyte activation also promotes freezing in zebrafish, consistent with our observed reduced exploratory behavior and NE-mediated passivity in larval zebrafish via hindbrain radial astrocytes (Mu et al., 2019).

Increased noise correlation in neuronal visual responses is suggested to be important for stimulus detection at the expense of fine discrimination (Azeredo da Silveira and Rieke, 2021).

Consistently, astrocyte activation by NE significantly increased network correlation, favoring detection over detailed encoding, as expected with perceived threats. Noradrenergic signaling has been associated with enhanced detection and rapid learning in certain brain areas, such as fear conditioning in the amygdala in adult mice (Uematsu et al., 2017). A similar role in highly plastic, developing circuits, could potentially bias the developmental refinement of circuits to optimize processing of specific environmental stimuli in a experience-dependent manner, filtered by neuromodulation. Future experiments in the developing animal should assess how differential circuit remodeling may occur under basal versus NE and astrocyte-regulated brain states.

While our investigations focused on the role of NE in the activation of radial astrocytes and the modulation of visual response properties in the OT, multiple other important neuromodulators have also been shown to modulate the function of the OT in a number of different vertebrate species. In zebrafish, 5-HT has been shown to regulate sensory processing through modulation of tectal circuits (Yokogawa et al., 2012) and also induce CNS wide alterations in network activity during shifts between exploratory swimming behavior (Marques et al., 2020). While we did not find acute effects of 5-HT on visual processing in the Xenopus laevis OT, it does have chronic effects on tectal function (Udoh et al., 2024). RGC axons and local interneurons are also known to express acetylcholine receptors in the OT and play an important role in tectal processing, including the modulation of inhibition and visual gain (Henley et al., 1986; Baginskas et al., 2013; Weigel and Luksch, 2014; Baginskas and Kuras, 2016; Asadollahi and Knudsen, 2016; Marques et al., 2020). Dopamine has long been known to alter visual processing (see review by Woolrych et al., 2021) and has been shown to alter visually-driven motor activity through modulation of the OT (Pérez-Fernández et al., 2017). How neuromodulators activate radial astrocytes in the OT, individually and in concert with one another, and whether different neuromodulators exert distinct effects on astrocytes and the function of the OT will be important avenues to explore.

## Methods

### Resource Availability

All data included in the publication of this manuscript will be made accessible by the lead contact upon reasonable request and have been uploaded to the Figshare repository (project 255941). All code written in house for this manuscript has been deposited on GitHub (https://github.com/nbenfey/Norepinephrine-Paper-Python-Scripts) and is publicly accessible as of the publication of this manuscript. Any additional requests for information relevant for the analysis of the data provided in this manuscript will be made available from the lead contact upon request. Any reagents produced for the experiments carried out in this manuscript will be made accessible upon request to the lead contact or through the relevant public repository responsible for distributing the reagents.

### Lead Contact

All requests for further information about resources or reagents used in this manuscript can be directed to the lead contact, Edward S. Ruthazer.

### Experimental model and subject details

All experimentation and use of animals was carried out in accordance with the Canadian Council of Animal Care guidelines and received approval from the Montreal Neurological Institute-Hospital Animal Care Committee. All adult Xenopus laevis frogs (albino) used to produce in vitro born tadpoles were kept in the care of the Montreal Neurological Institute-Hospital Center for Neurological Disease Models animal care facility. All animals used in the experiments outlined in this manuscript ranged from developmental stage 45 to stage 48. Tadpoles were reared in 0.1X Modified Barth’s Solution with HEPES (MBSH). As sexual characteristics are not defined at these early developmental stages, the sex of the animals used in our experiments was not relevant to assess.

### Tyrosine hydroxylase immunohistochemistry

St 48 tadpoles were fixed in 4% Paraformaldehyde in 0.2M Phosphate Buffer (EMS cat# 15735-30S) overnight at 4C. Tadpoles were cryoprotected sequentially in 15% fish gelatin (Norland) / 15% sucrose and 25% fish gelatin/ 15% sucrose overnight. Animals were then embedded in 20% fish gelatin/ 15% sucrose and cryosectioned horizontally with a Leica cryostat. 14µm sections were collected on Superfrost Plus slides (Fisher) and then stained with chicken anti-TH antibody. Briefly, sections were incubated with 1% sodium dodecyl sulfate for 5 min and then blocked in 5% normal goat serum and 1% bovine serum albumin in Tris buffered saline. Sections were stained with chicken anti-Tyrosine Hydroxylase (TH) antibody (Aves labs, TYH, 1:500, RRID: AB_10013440) and TH positive cells visualized with goat anti-chicken Alexa 488 (Thermo Fisher, A-11039, RRID: AB_2534096), nuclei were labelled with Dapi. Images were then acquired with a Zeiss Axio Imager M1.

### GCaMP6s mRNA synthesis and generation of GCaMP6s-expressing tadpoles

GCaMP6s-expressing tadpoles were generated following previously published methods from our lab (see Benfey et al., 2021 for more detail). In brief GCaMP6s mRNA was generated using a SP6 mMessage mMachine Kit (Ambion, Thermo Fisher) to transcribe GCaMP6s mRNA from pCS2+ plasmids. Following in vitro fertilization, eggs were monitored until the first cell division commenced and then injected with 500-750 pg of GCaMP6s mRNA using a calibrated glass micropipette and a PLI-100 picoinjector (Harvard Apparatus). Animals were raised under a 12 H light / 12 H dark cycle in an 18°C incubator until they reached stages developmental 46-48.

### Tectal electroporation of radial astrocytes

To transfect radial astrocytes with either membrane-targeted eGFPf for structural visualization or mTRPV1red for chemogenetic activation we followed previously published protocols from our lab (Tremblay et al., 2009). In brief, the tectum of anesthetized developmental stage 45 tadpoles (0.02% MS-222 in 0.1X MBSH) was injected with plasmid (0.5-1 μg/μL) encoding the desired construct and using two platinum electrodes placed on either side of the tectum current was passed in both directions of polarity using Grass Instruments SD9 electrical stimulator. Animals were left to recover in 0.1X MBSH for 2 days before chemogenetic activation and imaging was performed.

### Preparing GCaMP6s-expressing tadpoles for live imaging

After screening stage 46-48 GCaMP6s-expressing tadpoles for sufficient levels of GCaMP6s expression in the optic tectum using an Olympus BX-43 epifluorescence microscope, animals were treated with 2mM pancuronium dibromide (Tocris) in 0.1X MBSH for several minutes until the onset of paralysis and then embedded in 0.8% low melting point agarose (Thermo Fisher) in a custom-made imaging chamber. To circumvent the blood brain barrier and allow for direct acute pharmacological manipulation of the optic tectum during live imaging, a small puncture through the skin covering the tectum was cut using the tip of a 30 ga syringe needle 30 min before imaging. Animals were then submerged in 9mL of external saline solution (115mM NaCl, 2mM KCl, 3mM CaCl2, 3mM MgCl2, 5mM HEPES, and 10mM glucose, pH 7.20, 250 mOsm).

### Two-photon microscope

All in vivo imaging experiments were carried out using a Thorlabs multiphoton resonant scanner microscope with a 20X water-immersion objective (1.0 NA) attached to a piezoelectric focus mount (PI). A Spectra-Physics InSightX3 infrared femtosecond pulsed laser was used to generate fluorescence excitation for live imaging experiments.

### In vivo calcium imaging of GCaMP6s-expressing tadpoles

To minimize the occurrence of drift across all three dimensions during imaging sessions, animals were left to settle in the imaging chambers under the microscope for approximately 30 min before the onset of imaging. Calcium imaging was then performed in 5 min intervals using an excitation wavelength of 910 nm and a power of ∼ 125 mW (as measured before the scan-head). Each 5 min imaging period involved the collection of 4500 images at a rate of 15 images per second in a single optical section in the z dimension with x,y dimensions of 224.256 μm squared at a resolution of 512 pixels squared. Using ImageJ, images were then saved as tiff stacks for further processing.

All drugs used for pharmacological manipulation of the tectum were dissolved in 1 mL of external saline solution to facilitate rapid diffusion when applied. Agonists were applied immediately following collection of a 5 min baseline and their effects were imaged after waiting 60 s for water soluble drugs such as NE and after 300 s for less water-soluble drugs like capsaicin in order to allow for adequate diffusion of the agents into the optic tectum. Antagonists were added to the bath of embedded animals immediately following the creation of the incision above the tectum and allowed to equilibrate for the 30 min, the animals were left to settle prior to imaging.

Visual stimulation during live imaging experiments was carried out by using a flat panel display screen (800 × 480, Adafruit, NY) covered with a red filter (Wratten #29, Kodak) to stimulate one eye of the animal while imaging green fluorescence from the contralateral optic tectum.

### Electrophysiological experiments

Tectal whole-cell patch clamp recordings were made from the isolated whole brains of stage 46-48 albino *Xenopus laevis* tadpoles. Tadpoles were anesthetized in MS-222 (0.02% w/v in 0.1X MBSH, Sigma) and pinned to a Sylgard block submerged in ice chilled extracellular saline (115 mM NaCl, 4 mM KCl, 3 mM CaCl2, 3 mM MgCl2, 5 mM HEPES, and 10 mM glucose (all Sigma), pH 7.2 adjusted with NaOH, osmolarity: 250 mOsm) for surgery. An incision was made along the dorsal midline of the optic tectum to expose the ventricular surface and the brain was dissected out and pinned to a Sylgard block in a recording chamber filled with room temperature extracellular saline. When required, tetrodotoxin (TTX; 1 µM, Alomone Labs), picrotoxin (PTX; 100 µM, Abcam), 2,3-dihydroxy-6-nitro-7-sulfamoyl-benzo[f]quinoxaline-2,3-dione (NBQX; 20 µM, Tocris), and 2-amino-5-phosphonovaleric acid (D-APV; 50 µM, Tocris) were included in the extracellular saline. Using a glass micropipette (Sutter Instruments) with a broken tip, sections of the ventricular membrane were carefully removed to gain access to the tectal neuron cell bodies below. Individual tectal neurons were visualized using an Olympus BX51 upright microscope with a LUMPLFLN 60X (1.0 NA) water-immersion objective and a CCD camera (Sony XC-75).

Whole-cell voltage clamp recordings were performed at room temperature using glass micropipettes (6-10 MΩ) filled with cesium based internal saline (90 mM CsMeSO4, 5 mM MgCl2, 20 mM tetraethylammonium (TEA), 10 mM ethylene glyco-bis (β-aminoethylether)-N,N,N’,N’-tetraacetic acid (EGTA), 20 mM HEPES, 2 mM ATP, 0.3 mM GTP (all Sigma), pH 7.2 adjusted with CsOH, osmolarity 250 mOsm). Upon entering whole cell mode, the intracellular saline and cytoplasm were given 5 min to equilibrate before any recordings were made. Electrical signals were measured using a Multiclamp 700B amplifier, digitized at 10 KHz (Digidata 1440A) and recorded using pClamp 10.3 (Molecular Devices). Recordings were restricted to the center of the tectum to avoid any developmental variabilities along the rostrocaudal axis (Wu et al., 1996). mEPSCs were recorded at −70 mV in the presence of TTX and PTX and mIPSCs at 0 mV in the presence of TTX, NBQX, and APV. For paired pulse recordings, a 25 m cluster stimulating electrode (FHC Inc.) was placed on the optic chiasm to deliver a pair of 100 s pulses (50 ms ISI) to retinal ganglion cell axons from an ISO-flex stimulation isolation unit (AMPI). Paired pulse recordings were made at −70 mV in the presence of PTX. In whole cell-current clamp experiments, a potassium gluconate based internal saline (100 mM K gluconate, 8 mM KCl, 5 mM NaCl, 1.5 mM MgCl2, 20 mM HEPES, 10 mM EGTA, 2 mM ATP, 0.3 mM GTP (all Sigma), pH 7.2 adjusted with KOH, osmolarity 250 mOsm) was used to permit spiking. A constant current, set at the beginning of the experiment to bring the membrane potential to −65 mV, was injected throughout.

All electrophysiological data were analyzed using Clampfit 11.2. mEPSCs and mIPSCs were identified automatically using custom detection templates in Clampfit 11.2 and all events were verified to remove false positives. The mean frequency and amplitude of mPSCs were measured in 5-min-long traces starting 5 min after drug application and normalized to a baseline trace.

Paired pulse ratios were calculated by dividing the peak amplitude of the second evoked PSC by the first. In current clamp experiments, deviations in membrane potential from −65 mV following drug application were measured in 30 s bins. Intrinsic excitability was calculated as the number of spikes evoked by 20 pA steps of injected depolarizing current over a range of 0-100 pA (200 ms square pulses, 20 s ISI) before and 5 min after drug application.

### Visual stimulation paradigms

Three different visual stimulation paradigms were used throughout our experiments. All visual stimuli were generated using custom python scripts running the PsychoPy package (https://www.psychopy.org/index.html). All stimuli were presented with an interstimulus interval of 20 s to avoid the habituation that we have observed to occur at faster stimulus presentation frequencies. Looming stimuli consisted of a blank screen for 19 s followed by the appearance of a small black dot which rapidly expands to fill the screen over the following 1 s. The moving dots stimuli used in the moving dots vs loom experiments were presented for 1 s following 19 s of blank screen and consisted of 25 size 25 black dots (∼10° in size for the animal) which randomly appear on the screen and move at a speed of 2 units per frame in various directions with no coherence. The moving dots used in the zero coherence versus full coherence experiments persisted for 2 s to exaggerate the motion of the stimulus and again consisted of 25 size 25 black dots (∼10°) which randomly appear on the screen and move at a speed of 2 units per frame either with no coherence or with full coherence in a downward direction to enhance visual detection.

### Registration and ROI extraction of raw calcium imaging data using Suite2p

Calcium imaging data collected on the two-photon microscope was registered and both neuronal and radial astrocyte ROIs were extracted using Suite2p software (http://github.com/MouseLand/suite2p) running on python 3 as described previously (Benfey et al., 2021). Raw fluorescence traces for each ROI were exported for further analysis.

### Baseline normalization and quantification of fluorescence increases

Raw fluorescence traces were normalized to baseline relying on standard ΔF/F0 methods. In brief, baseline fluorescence was estimated using a sliding window to calculate the bottom 10^th^ percentile of fluorescence values within a 100 s window centered around each value in the fluorescence trace. For each fluorescence value, we subtracted this baseline and then divided by the baseline value + 10 in order to obtain a percent change in the fluorescence signal relative to baseline across time. The addition of 10 to the baseline value before division leads to an underestimation of true ΔF/F0 but prevents artifacts that arise due to divisions of baselines close to 0.

### Correlation and principal component analysis

Correlation and principal component analysis of the tectal network activity during each 5 min imaging epoch was computed using custom made python scripts built using open source modules from both scikit-learn (https://scikit-learn.org/stable/index.html) and SciPy (https://scipy.org/) which we have made available along with this manuscript.

### Exploratory and Loom-evoked escape behavior

Loom-evoked escape behavior was assayed in freely swimming stage 47 tadpoles using methods published by Lim and Ruthazer (2021)(https://github.com/tonykylim/XenLoom_beta). A small incision was made over the brain ventricle to expose the brain to capsaicin (10 µM) in rearing solution for 5 min prior to testing. The tadpole was then transferred in a closed 6 cm diameter petri dish submerged in 0.1X MBSH at the bottom of a large 20 cm glass bowl filled with 0.1X MBSH and allowed to freely navigate this environment. A 2000 lumen projector was used to present 10 looming stimuli (one every 20 s) on a paper screen attached to the side of the bowl while a webcam fixed overtop of the bowl tracked the animal’s movement through its environment. Custom python scripts published by Lim and Ruthazer (2021) utilizing computer vision to identify and track tadpole motion were used to assess loom-evoked escape responses while blind to experimental conditions. Animals that swam a distance greater than 10 percent of the width of the petri dish during the inter-looming-stimulus interval were considered to be exhibiting exploratory behaviour during that time interval.

### Statistical analyses

All statistical analysis was performed using GraphPad’s Prism software package. All pairwise analysis was performed using paired or unpaired t tests as appropriate. Group analysis was performed using one-way or two-way repeated measures ANOVAs followed by Holm-Šidák’s multiple comparisons tests as appropriate. All relevant statistical information including p values (with significance defined as a p value of 0.05 or less) can be found in the appropriate supplementary tables.

## Acknowledgements

We would like to thank the lab of DA Prober for providing us with the mTRPV1red plasmid we used for our chemogenetic experiments. This research was supported by both Masters and Doctoral Canada Postgraduate Scholarships to N.J.B from the Canadian Institutes of Health Research and the Natural Sciences and Engineering Research Council of Canada. Support was also received through the funding E.S.R. received from the Canadian Institutes of Health Research (FDN-143238; PJT-180478) and Fonds de recherche santé Québec (31036). E.S.R also received financial support from the Brain Canada Canadian Neurophotonics Platform.

## Author Contributions

N.J.B and E.S.R conceived of and designed the study. N.J.B. carried out all imaging experiments, except for the 5-HT dataset which was collected by E.C, and the mTRPV1red and SCH 58261 datasets which were collected by D.F. Data analysis of the imaging data was carried with the help of custom python scripts written by N.J.B and A.T.B. F.C. collected and analyzed the electrophysiological data for this study. A.S. developed the methodology for labeling animals with GCaMP6s and carried out the immunohistochemistry as well as provided technical support throughout the development of the techniques used in this study. N.J.B. and O.R. collected and analyzed the behavioral data for this study. N.J.B. drafted the manuscript which was subsequently edited by E.S.R. All authors contributed to the editing of the final version of the manuscript. E.S.R obtained the funding and supervised the research.

## Declaration of Interests

The authors declare no competing interests.

## Notes

https://figshare.com/projects/Norepinephrine_acts_through_radial_astrocytes_in_the_developing_optic_tectum_to_enhance_threat_detection_and_escape_behavior/255941

https://github.com/nbenfey/Norepinephrine-Paper-Python-Scripts

### Competing Interest Statement

The authors have declared no competing interest.

