## Supplemental materials for "Norepinephrine acts through radial astrocytes in the developing optic tectum to enhance threat detection and escape behavior"

**Figure S1. Additional measures related to the effects of norepinephrine on the activity of neurons in the optic tectum.**

**(A)** Retinotectal PPR is unaltered by vehicle wash-on. **(B)** NE application has no effect on tectal neuron resting membrane potential. (**C**) NE does not change the mean number of spikes produced by current injections in tectal neurons. **(D)** Representative traces of miniature inhibitory postsynaptic currents (mIPSCs) recorded at 0 mV with glutamatergic transmission pharmacologically blocked in tectal neurons during baseline (*top*) and following addition of NE (*bottom*). **(E, F)** No changes were observed in mIPSC frequency in vehicle treated (E) and NE treated (F) animals. **(G, H)** mIPSC amplitude in vehicle (G) and NE treated (H) animals was also unaltered.

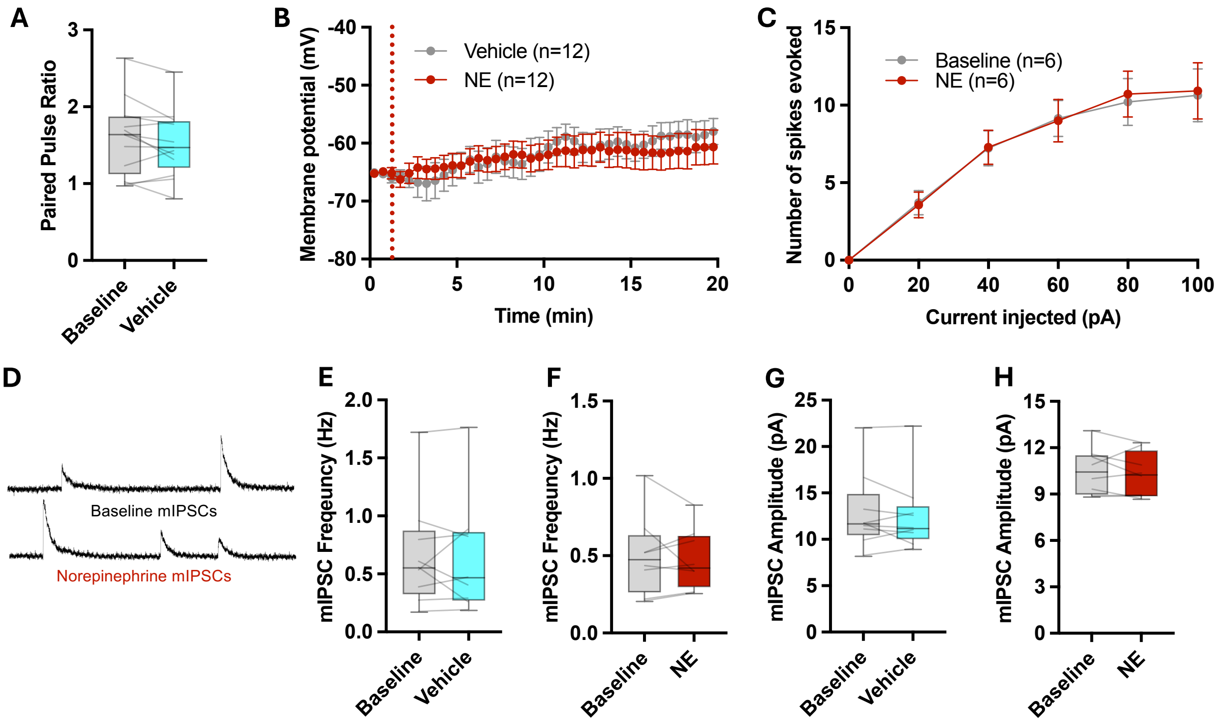


Benfey et al., Figure S1

**Figure S2. Norepinephrine enhances the detection of coherent motion in the optic tectum.**

**(A)** Baseline normalized ($\Delta F/F_0$) traces of the calcium activity of radial astrocytes and tectal neurons in a single optical section of one hemisphere of the OT during visual stimulation with alternating stimuli of small moving dots with no coherent motion and small moving dots with full coherent motion under baseline conditions (*left*) and following addition of NE (100 μM) (*right*). **(B, C)** Quantification of the number of responsive radial astrocytes (B) and tectal neurons (C) in a single optical section of the OT exhibiting calcium events over the course of 5 min of imaging. **(D)** Quantification of the average Pearson correlation coefficient between tectal neurons during baseline and NE conditions. **(E)** Quantification of the percentage of the total variance captured by PC1 in baseline and NE conditions. **(F)** Average peak response amplitudes of tectal neurons to presentation of dots moving in random directions with no coherence (C=0) (*left*) and fully coherently moving dots (C=1) (*right*). **(G)** Quantification of the mean peak response amplitudes to moving dots with no coherent motion (*left*) and moving dots with full coherent motion (*right*) of tectal neurons during baseline and NE conditions. **(H)** Representative traces showing the average response of tectal neurons to both stimuli in each coherence preference ratio bin. **(I)** Histogram of the coherent motion preference ratio of tectal neurons during baseline and NE conditions and **(J)** mean neuronal coherent motion preference per animal during baseline and NE conditions both show a shift favoring coherent motion. See Table S2 for statistical details.

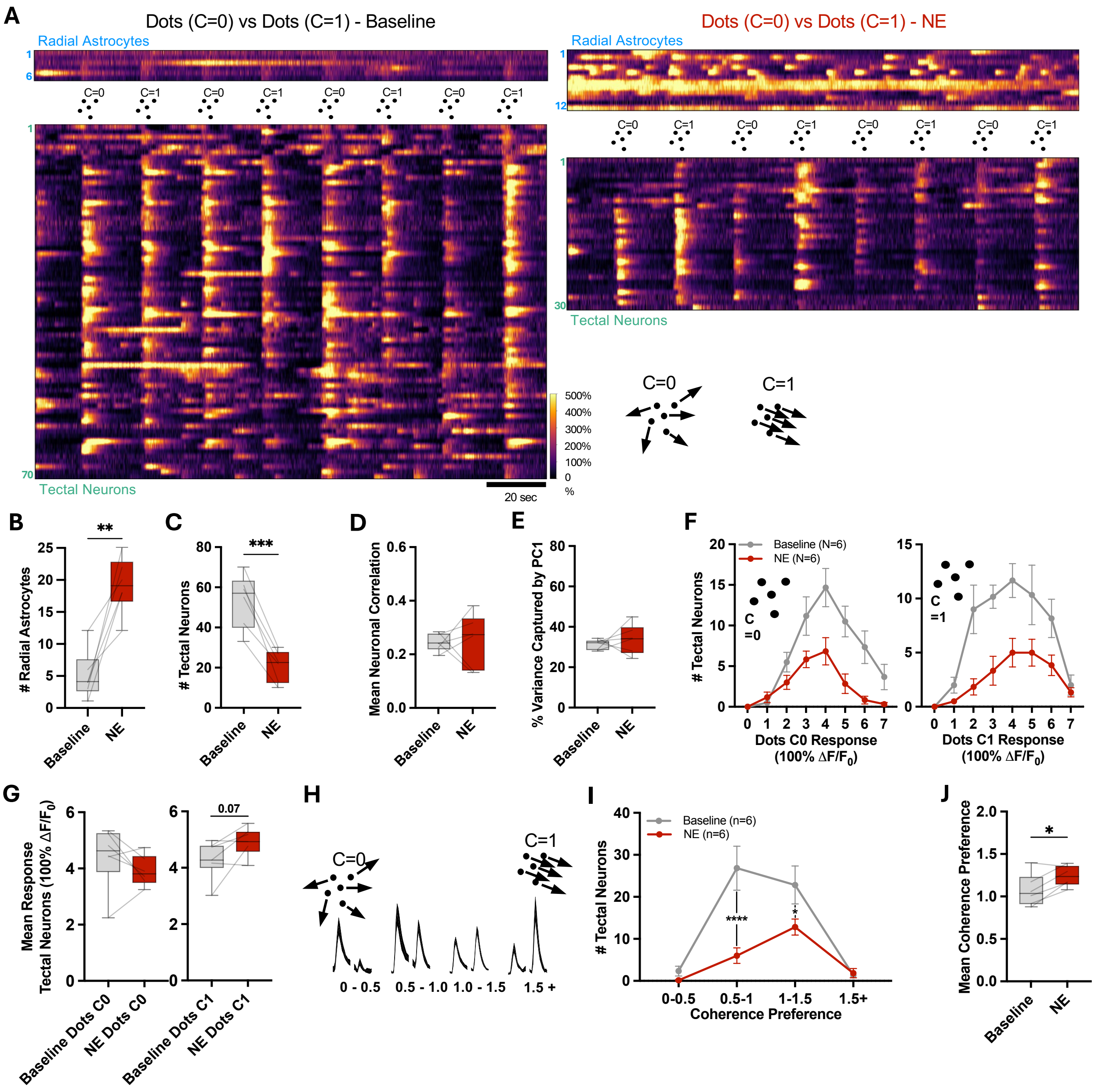


Benfey et al., Figure S2

**Figure S3. Absence of effects of serotonin on visually driven neuronal activity in the optic tectum.**

**(A)** Baseline normalized ($\Delta F/F_0$) traces of the calcium activity of radial astrocytes and tectal neurons in a single optical section of one hemisphere of the OT during visual stimulation with alternating stimuli of moving dots and looms following the application of serotonin (5-HT, 100 µM). **(B, C)** Quantification of the number of radial astrocytes (B) and tectal neurons (C) in a single optical section of the OT exhibiting calcium events over the course of 5 min of imaging. **(D)** Average Pearson correlation coefficient between tectal neurons per animal. **(E)** Fraction of variance captured by PC1 in each condition. **(F)** Histogram of peak response amplitudes to moving dots (*left*) and looming (*right*) stimuli in tectal neurons. **(G)** Mean peak response amplitudes to moving dots (*left*) and looming (*right*) stimuli of tectal neurons per animal. **(H)** Loom preference ratio of tectal neurons and **(I)** mean loom preference of tectal neurons per animal. Note that 5-HT produced no significant effects compared to baseline measurements.

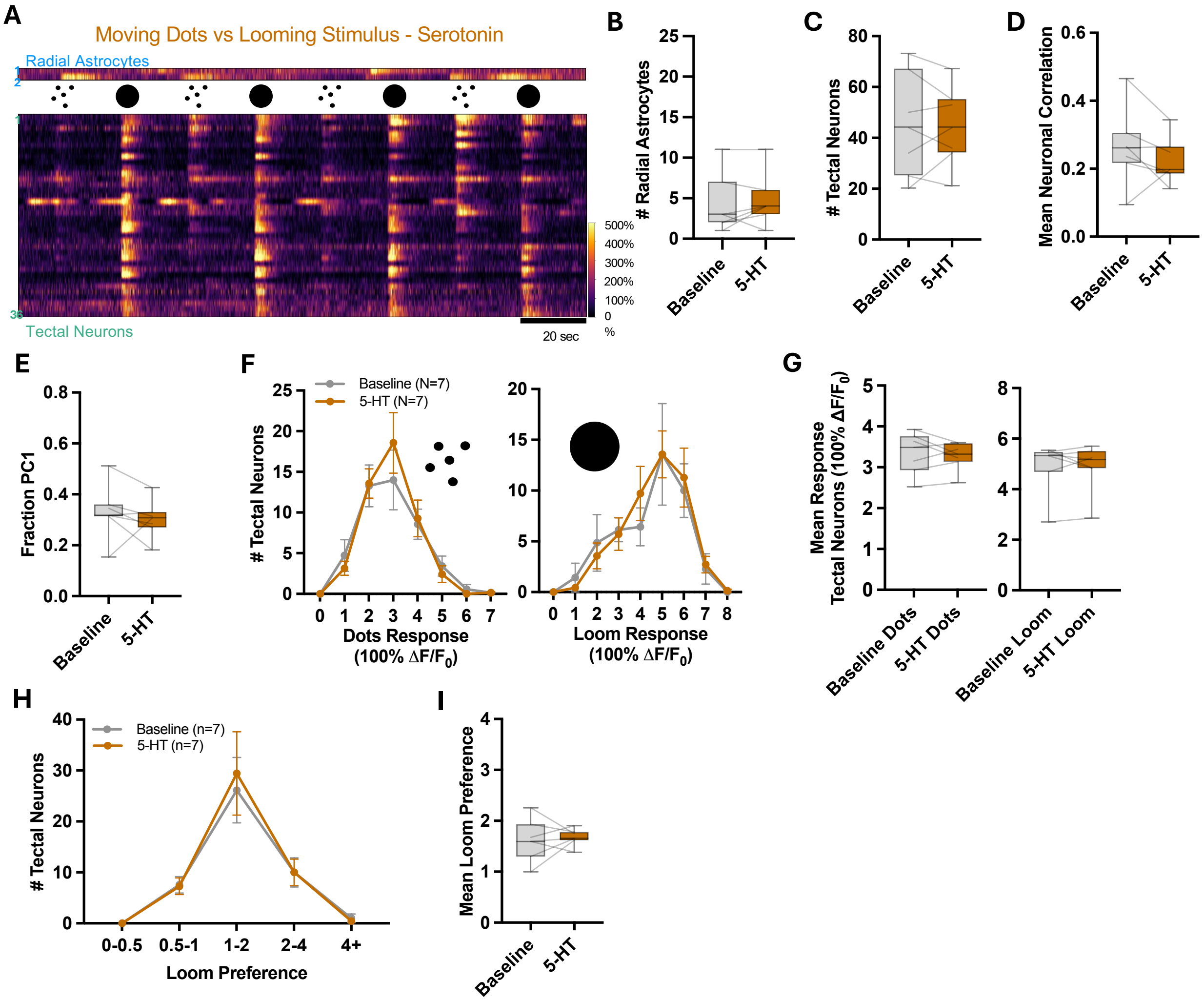


Benfey et al., Figure S3

**Figure S4. Blockade of adenosine type 1 receptors or hemichannels also modulates the tectal response to norepinephrine.**

**(A)** Baseline normalized ($\Delta F/F_0$) traces of the calcium activity of radial astrocytes (*top*) and tectal neurons (*bottom*) in a single optical section of one hemisphere of the OT during visual stimulation with alternating stimuli of small moving dots and loom under adenosine A1 receptor blockade with DPCPX (100 nM) followed by the addition of NE (100 μM) (*left*) and gap junction/hemichannel inhibition with carbenoxolone (CBX) (100 μM), followed by the addition of NE (100μM) (*right*). **(B)** Number of radial astrocytes active during the recordings in each condition. **(C)** Number of tectal neurons active during the recordings in each condition. **(D)** Mean correlation between tectal neurons active during the recordings in each condition. **(E)** Percentage of the total variance captured by PC1 in each condition (Table S4). **(F)** Average peak response amplitudes to moving dots (*left*) and looming (*right*) stimuli in tectal neurons in each condition. n=8 animals NE treated, n=6 animals DPCPX+NE treated, n=7 animals CBX+NE treated. **(G, H)** Quantification of the loom preference ratio of tectal neurons in DPCPX, DPCPX + NE wash-on, and NE wash-on conditions (G) and CBX, CBX + NE, and NE treated conditions (H). **(I)** Mean loom preference of tectal neurons during each condition See Table S4 for statistical details.

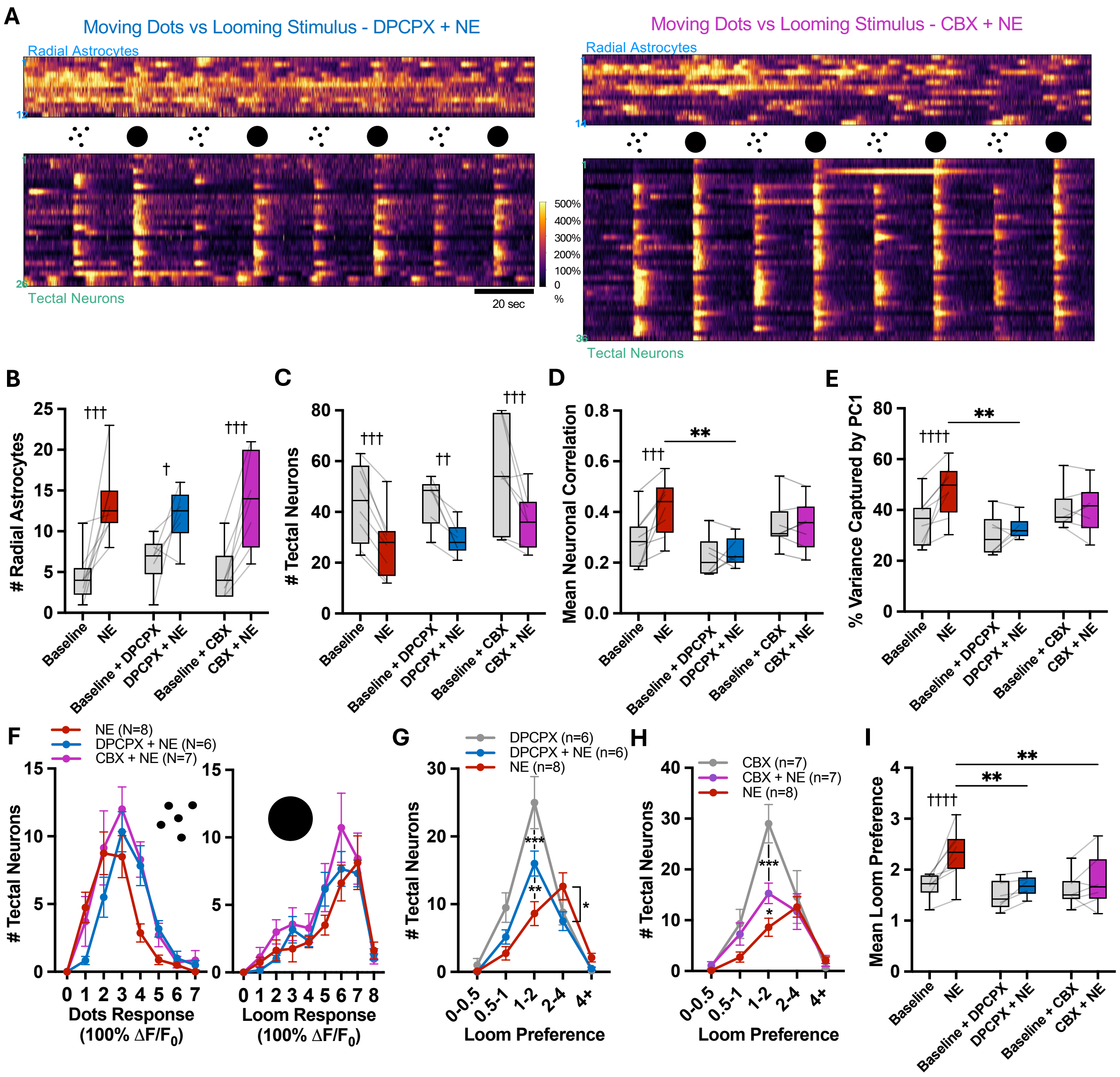


Benfey et al., Figure S4

**Video legends**

**Video S1. Norepinephrine elevates calcium in radial astrocytes but not neurons when neuronal spiking activity is silenced.** GCaMP6s z-projection through 150 μm depth in the right OT hemisphere after TTX (1μM) wash-on (left), and subsequent NE (100 μM) application (right).

**Video S2. Norepinephrine elevates the spontaneous activity of radial astrocytes and decreases the activity of tectal neurons during resting state.** Average GCaMP6s signal in a single optical section during baseline (left) and after NE (100 μM) (right).

**Video S3. Norepinephrine selectively alters visual processing in the optic tectum.** Typical responses of the optic tectum to dots and loom stimuli during baseline (left) and after NE (100 μM) (right).

**Video S4. Tadpole demonstrating loom evoked escape behavior.** A freely swimming tadpole exhibits a characteristic escape response following visual presenation of a looming stimuli.

**Video S5. Tadpole demonstrating exploratory behavior.** A freely swimming tadpole exhibiting exploratory behavior in the periods of time between looming stimulus presentation.

**Supplemental Tables with statistical analyses for "Norepinephrine acts through radial astrocytes in the developing optic tectum to enhance threat detection and escape behavior" by Benfey et al.**

### Table S1

| Figure | Test Type | Sample Size | Statistics | Multiple Comparisons | Statistics |
|---|---|---|---|---|---|
| Figure 1G,H | Paired t-tests | (n=11 animals) | Glia: p=0.0001<br>Neurons p=0.0009 | | |
| Figure 1I | Two-Way ANOVA (Repeated-Measures) | (n=9 animals Vehicle)<br>(n=9animals NE) | $F_{Interaction}$=8.323 p=0.0108 | Holm-Sidak | Baseline vs Vehicle: p=0.7807<br>Baseline vs NE: p=0.0010 |
| Figure 1J | Unpaired t-test | (n=9 animals) | p=0.9405 | | |
| Figure 1K | Paired t-test | (n=15 animals) | p=0.0098 | | |
| Figure 1L | Unpaired t-test | (n=9 animals Vehicle)<br>(n=8 animals NE) | p=0.8834 | | |
| Figure 1M | Unpaired t-test | (n=9 animals Vehicle)<br>(n=8 animals NE) | p=0.9590 | | |
| Figure S1A | Paired t-test | (n=13 animals) | p=0.1061 | | |
| Figure S1B | Unpaired t-test | (n=12 animals in each group) | p=0.3611 | | |
| Figure S1C | Paired t-test | (n= 6 animals) | p=0.4009 | | |
| Figure S1E,F | Paired t-tests | (n=8 animals NE)<br>(n=9 animals Vehicle) | pNE=0.6870<br>pVehicle=0.8960 | | |
| Figure S1G,H | Paired t-tests | (n=8 animals NE)<br>(n=9 animals Vehicle) | pNE=0.4975<br>pVehicle=0.5077 | | |

### Table S2

| Figure | Test Type | Sample Size | Statistics | Multiple Comparisons | Statistics |
|---|---|---|---|---|---|
| Figure 2C,D | Paired t-tests | (n=8 animals) | Glia: p=0.0032<br>Neurons: p=0.0001 | | |
| Figure 2E | Paired t-test | (n=8 animals) | p=0.0072 | | |
| Figure 2F | Paired t-test | (n=8 animals) | p=0.0067 | | |
| Figure 2J | Pairted t-tests | (n=8 animals) | pDots=0.0031<br>pLoom=0.0457 | | |
| Figure 2K | Two-Way ANOVA (Repeated-Measures) | (n=8 animals) | $F_{Interaction}$=42.24<br>p<0.0001 | Holm-Sidak | Baseline vs NE:<br>LPR=0-0.5: p=0.6399<br>LPR=0.5-1: p<0.0001<br>LPR=1-2: p<0.0001<br>LPR=2-4: p=0.5464<br>LPR=4+: p=0.6399 |
| Figure 2L | Paired t-test | (n=8 animals) | p=0.0035 | | |
| Figure S2B,C | Paired t-tests | (n=6 animals) | Glia: p=0.0024<br>Neurons: p=0.0006 | | |
| Figure S2D | Paired t-tests | (n=6 animals) | p=0.8485 | | |
| Figure S2E | Paired t-tests | (n=6 animals) | p=0.5158 | | |
| Figure S2G | Paired t-tests | (n=6 animals) | pC0=0.2751<br>pC1=0.0711 | | |
| Figure S2I | Two-Way ANOVA (Repeated-Measures) | (n=6 animals) | $F_{Interaction}$=7.267<br>p=0.0017 | Holm-Sidak | Baseline vs NE:<br>CPR=0-0.5: p=0.7963<br>CPR=0.5-1: p<0.0001<br>CPR=1-1.5: p=0.0317<br>CPR=1.5+: p=0.8894 |
| Figure S2J | Paired t-test | (n=6 animals) | p=0.0171 | | |

### Table S3

| Figure | Test Type | Sample Size | Statistics | Multiple Comparisons | Statistics |
|---|---|---|---|---|---|
| Figure 3B | Two-Way ANOVA (Repeated-Measures) | (n=8 animals NE)<br>(n=7 animals PRZ+NE) | $F_{Interaction}$=8.138<br>p=0.0028 | Holm-Sidak | Baseline vs NE: p<0.0001<br>PRZ vs PRZ+NE: p=0.1831<br>Before Treatments:<br>NE vs PRZ+NE: p=0.7946<br>After Treatments:<br>NE vs PRZ+NE: p<0.0001 |
| Figure 3C | Two-Way ANOVA (Repeated-Measures) | (n=8 animals NE)<br>(n=7 animals PRZ+NE) | $F_{Interaction}$=7.075<br>p=0.0051 | Holm-Sidak | Baseline vs NE: p<0.0001<br>PRZ vs PRZ+N: p=0.2279<br>Before Treatments:<br>NE vs PRZ+NE: p=0.0809<br>After Treatments:<br>NE vs PRZ+NE: p=0.0048 |
| Figure 3D | Two-Way ANOVA (Repeated-Measures) | (n=8 animals NE)<br>(n=7 animals PRZ+NE) | $F_{Interaction}$=9.30<br>p=0.0015 | Holm-Sidak | Baseline vs NE: p=0.0003<br>PRZ vs PRZ+NE: p=0.4579<br>Before Treatments:<br>NE vs PRZ+NE: p=0.0580<br>After Treatments:<br>NE vs PRZ+NE: p=0.3818 |
| Figure 3E | Two-Way ANOVA (Repeated-Measures) | (n=8 animals NE)<br>(n=7 animals PRZ+NE)<br>(n=7 animals 5-HT) | $F_{Interaction}$=6.735<br>p=0.0062 | Holm-Sidak | Baseline vs NE: p=0.0004<br>PRZ vs PRZ+NE: p=0.5318<br>Before Treatments:<br>NE vs PRZ+NE: p=0.2792<br>After Treatments:<br>NE vs PRZ+NE: p=0.4670 |
| Figure 3G | Two-Way ANOVA (Repeated-Measures) | (n=7 animals PRZ+NE)<br>(n=8 animals NE) | $F_{Interaction}$=2.955<br>p=0.0054 | Holm-Sidak | PRZ vs PRZ+NE:<br>LPR=0-0.5: p=0.9948<br>LPR=0.5-1: p=0.6352<br>LPR=1-2: p=0.5394<br>LPR=2-4: p=0.4263<br>LPR=4+: p=0.9819<br>PRZ+NE vs NE:<br>LPR=0-0.5: p=0.9948<br>LPR=0.5-1: p=0.2439<br>LPR=1-2: p<0.0001<br>LPR=2-4: p=0.4263<br>LPR=4+: p=0.9819 |

| | | | | | |
|---|---|---|---|---|---|
| Figure 3H | Two-Way ANOVA (Repeated-Measures) | (n=8 animals NE)<br>(n=7 animals PRZ+NE) | $F_{Interaction}$=5.512<br>p=0.0129 | Holm-Sidak | Baseline vs NE: p=0.0004l<br>PRZ vs PRZ+NE: p=0.9130<br>Before Treatments:<br>NE vs PRZ+NE: p=0.7961<br>After Treatments:<br>NE vs PRZ+NE: p=0.0104 |
| Figure 3J | One-Way ANOVA (Repeated-Measures) | (n=5 animals) | F=11.91<br>p=0.0056 | Holm-Sidak | Baseline vs YOH: p=0.0488<br>YOH+PRZ vs YOH+PRZ+NE: p=0.7040 |
| Figure 3K | One-Way ANOVA (Repeated-Measures) | (n=9 animals Vehicle)<br>(n=9 animals NE)<br>(n=8 animals PRZ+NE) | F=0.4199<br>p=0.0005 | Holm-Sidak | Vehicle vs NE: p=0.0011<br>Vehicle vs PRZ+NE: p=0.7974<br>NE vs PRZ+NE: p=0.0019 |
| Figure 3K | One-Way ANOVA (Repeated-Measures) | (n=9 animals Vehicle)<br>(n=9 animals NE)<br>(n=8 animals PRZ+NE) | F=0.5316<br>p=0.5947 | | |
| Figure 3L | Paired t-tests | (n=8 animals) | pFrequency=0.0446<br>pAmplitude=0.0926 | | |
| Figure S3B,C | Paired t-tests | (n=7 animals) | Glia: p=0.4128<br>Neurons p=0.9107 | | |
| Figure S3D | Paired t-test | (n=7 animals) | p=0.2327 | | |
| Figure S3E | Paired t-test | (n=7 animals) | p=0.4339 | | |
| Figure S3G | Paired t-tests | (n=7 animals) | pDots=0.9836<br>pLoom=0.5023 | | |
| Figure S3H | Two-Way ANOVA (Repeated-Measures) | (n=7 animals 5-HT) | $F_{Interaction}$=0.7204<br>p=0.5848 | | |
| Figure S3I | Paired t-test | (n=7 animals) | p=0.7079 | | |

### Table S4

| Figure | Test Type | Sample Size | Statistics | Multiple Comparisons | Statistics |
|---|---|---|---|---|---|
| Figure 4C | Paired t-test | (n=8 animals) | pFrequency=0.0001 | | |
| Figure 4E | Paired t-test | (n=13 animals) | p=0.0174 | | |
| Figure 4F | One-Way ANOVA (Repeated-Measures) | (n=9 animals Vehicle)<br>(n=9 animals NE)<br>(n=9 animals AMPCP+NE)<br>(n=10 animals DPCPX+NE)<br>(n= 9 animals SCH 58261+NE)<br>(n=9 animals MRS 1754+NE) | F=7.226<br>p<0.0001 | Holm-Sidak | Vehicle vs NE: p=0.0011<br>Vehicle vs AMPCP+NE: p=0.1650<br>Vehicle vs DPCPX+NE: p=0.0008<br>Vehicle vs SCH 58261+NE: p=0.8843<br>Vehicle vs MRS 1754+NE: p=0.0014 |
| Figure 4G | One-Way ANOVA (Repeated-Measures) | (n=9 animals Vehicle)<br>(n=9 animals NE)<br>(n=9 animals AMPCP+NE)<br>(n=10 animals DPCPX+NE)<br>(n= 9 animals SCH 58261+NE)<br>(n=9 animals MRS 1754+NE) | F=2.571<br>p=0.0383 | Holm-Sidak | Vehicle vs NE: p=0.9970<br>Vehicle vs AMPCP+NE: p=0.7566<br>Vehicle vs DPCPX+NE: p=0.9970<br>Vehicle vs SCH 58261+NE: p=0.1699<br>Vehicle vs MRS 1754+NE: p=0.4386 |
| Figure 4H | Two-Way ANOVA (Repeated-Measures) | (n=8 animals NE)<br>(n=9 animals SCH 58261+NE) | $F_{Interaction}$=2.059<br>p=0.1718 | Uncorrected Fisher's LSD | Baseline vs NE: p=0.0001<br>SCH 58261 vs SCH 58261+NE: p=0.0053 |
| Figure 4I | Two-Way ANOVA (Repeated-Measures) | (n=8 animals NE)<br>(n=9 animals SCH 58261+NE) | $F_{Interaction}$=23.39<br>p=0.0002 | Uncorrected Fisher's LSD | Baseline vs NE: p<0.0001<br>SCH 58261 vs SCH 58261+NE: p=0.2430<br>Baseline vs SCH 58261: p<0.0001<br>NE vs SCH 58261+NE: p=0.0019 |
| Figure 4J | Two-Way ANOVA (Repeated-Measures) | (n=8 animals NE)<br>(n=9 animals SCH 58261+NE) | $F_{Interaction}$=23.57<br>p=0.0002 | Uncorrected Fisher's LSD | Baseline vs NE: p=0.0003<br>SCH 58261 vs SCH 58261+NE: p=0.0501<br>Baseline vs SCH 58261: p=0.0857<br>NE vs SCH 58261+NE: p<0.0001 |
| Figure 4K | Two-Way ANOVA (Repeated-Measures) | (n=8 animals NE) | $F_{Interaction}$=14.00<br>p=0.0020 | Uncorrected Fisher's LSD | Baseline vs NE: p=0.0005<br>SCH 58261 vs SCH 58261+NE: p=0.4349 |

| | | | | | |
|---|---|---|---|---|---|
| | | (n=9 animals SCH 58261+NE) | | | Baseline vs SCH 58261: p=0.4340<br>NE vs SCH 58261+NE: p=0.0002 |
| Figure 4M | Two-Way ANOVA (Repeated-Measures) | (n=8 animals NE)<br>(n=9 animals SCH 58261+NE) | $F_{Interaction}$=7.089<br>p<0.0001 | Holm-Sidak | SCH 58261 vs SCH 58261+NE:<br>LPR=0-0.5: p>0.9999<br>LPR=0.5-1: p=0.7947<br>LPR=1-2: p=0.4620<br>LPR=2-4: p=0.4620<br>LPR=4+: p=0.7824<br>SCH 58261+NE vs NE:<br>LPR=0-0.5: p=0.9936<br>LPR=0.5-1: p=0.7947<br>LPR=1-2: p=0.0719<br>LPR=2-4: p<0.0001<br>LPR=4+: p=0.2397 |
| Figure 4N | Two-Way ANOVA (Repeated-Measures) | (n=8 animals NE)<br>(n=9 animals SCH 58261+NE) | $F_{Interaction}$=15.35<br>p=0.0014 | Uncorrected Fisher's LSD | Baseline vs NE: p=0.0022<br>SCH 58261 vs SCH 58261+NE: p=0.0928<br>Baseline vs SCH 58261: p=0.3967<br>NE vs SCH 58261+NE: p=0.0004 |
| Figure S4B | Two-Way ANOVA (Repeated-Measures) | (n=8 animals NE)<br>(n=6 animals DPCPX+NE)<br>(n=7 animals CBX+NE) | $F_{Interaction}$=0.9400<br>p=0.4090<br>$F_{Treatment}$=45.21<br>p<0.0001 | Holm-Sidak | Baseline vs NE: p=0.0001<br>DPCPX vs DPCPX+NE: p=0.0204<br>CBX vs CBX+NE: p=0.0003 |
| Figure S4C | Two-Way ANOVA (Repeated-Measures) | (n=8 animals NE)<br>(n=6 animals DPCPX+NE)<br>(n=7 animals CBX+NE) | $F_{Interaction}$=0.0377 p=0.9636<br>$F_{Treatment}$=42.09<br>p<0.0001 | Holm-Sidak | Baseline vs NE: p=0.0007<br>DPCPX vs DPCPX+NE: p=0.0042<br>CBX vs CBX+NE: p=0.0010 |
| Figure S4D | Two-Way ANOVA (Repeated-Measures) | (n=8 animals NE)<br>(n=6 animals DPCPX+NE)<br>(n=7 animals CBX+NE) | $F_{Interaction}$=6.793<br>p=0.0063 | Holm-Sidak | Baseline vs NE: p=0.0001<br>DPCPX vs DPCPX+NE: p=0.5831<br>CBX vs CBX+NE: p=0.8904<br>Before Treatments:<br>NE vs DPCPX+NE: p=0.3441<br>NE vs CBX+NE: p=0.3441<br>After Treatments:<br>NE vs DPCPX+NE: p=0.0027<br>NE vs CBX+NE: p=0.1495 |
| Figure S4E | Two-Way ANOVA (Repeated-Measures) | (n=8 animals NE)<br>(n=6 animals DPCPX+NE)<br>(n=7 animals CBX+NE) | $F_{Interaction}$=5.941<br>p=0.0104 | Holm-Sidak | Baseline vs NE: p=0.0002<br>DPCPX vs DPCPX+NE: p=0.3605<br>CBX vs CBX+NE: p=0.9581<br>Before Treatments: |

| | | | | | |
|---|---|---|---|---|---|
| | | | | | NE vs DPCPX+NE: p=0.4491<br>NE vs CBX+NE: p=0.4491<br>After Treatments:<br>NE vs DPCPX+NE: p=0.0061<br>NE vs CBX+NE: p=0.1027 |
| Figure S4G | Two-Way ANOVA (Repeated-Measures) | (n=6 animals DPCPX+NE)<br>(n=8 animals NE) | $F_{Interaction}$=7.210<br>p<0.0001 | Holm-Sidak | DPCPX vs DPCPX+NE:<br>LPR=0-0.5: p=0.8914<br>LPR=0.5-1: p=0.1308<br>LPR=1-2: p=0.0005<br>LPR=2-4: p=0.5706<br>LPR=4+: p=0.8871<br>DPCPX+NE vs NE:<br>LPR=0-0.5: p=0.9546<br>LPR=0.5-1: p=0.2730<br>LPR=1-2: p=0.0011<br>LPR=2-4: p=0.0428<br>LPR=4+: p=0.7086 |
| Figure S4H | Two-Way ANOVA (Repeated-Measures) | (n=7 animals CBX+NE)<br>(n=8 animals NE) | $F_{Interaction}$=4.058<br>p=0.0003 | Holm-Sidak | CBX vs CBX+NE:<br>LPR=0-0.5: p=0.9363<br>LPR=0.5-1: p=0.5406<br>LPR=1-2: p=0.0001<br>LPR=2-4: p=0.6187<br>LPR=4+: p=0.9063<br>CBX+NE vs NE:<br>LPR=0-0.5: p=0.9363<br>LPR=0.5-1: p=0.2835<br>LPR=1-2: p=0.0373<br>LPR=2-4: p=0.7733<br>LPR=4+: p=0.9685 |
| Figure S4I | Two-Way ANOVA (Repeated-Measures) | (n=8 animals NE)<br>(n=6 animals DPCPX+NE)<br>(n=7 animals CBX+NE) | $F_{Interaction}$=5.124<br>p=0.0173 | Holm-Sidak | Baseline vs NE: p<0.0001<br>DPCPX vs DPCPX+NE: p=0.1543<br>CBX vs CBX+NE: p=0.2370<br>Before Treatments:<br>NE vs DPCPX+NE: p=0.5478<br>NE vs CBX+NE: p=0.7225<br>After Treatments:<br>NE vs DPCPX+NE: p=0.0076<br>NE vs CBX+NE: p=0.0085 |

### Table S5

| Figure | Test Type | Sample Size | Statistics | Multiple Comparisons | Statistics |
|---|---|---|---|---|---|
| Figure 5C | Two-Way ANOVA (Repeated-Measures) | (n=6 wild type animals) (n=8 mTRPV1red transfected animals) | $F_{Capsaisin}$=10.57 p=0.0047 | Holm-Sidak | Wild Type Baseline vs Wild Type Capsaicin: p=0.2269<br>Transfected Baseline vs Transfected Capsaicin: p=0.0068 |
| Figure 5D | Unpaired t-test | (n= 6 wild type animals) (n=8 mTRPV1red transfected animals) | p=0.0809 | | |
| Figure 5F,G | Paired t-tests | (n=6 mTRPV1red transfected animals) | Glia: p=0.0023<br>Neurons: p=0.0063 | | |
| Figure 5H | Paired t-test | (n= 6 mTRPV1red transfected animals) | p=0.0576 | | |
| Figure 5I | Paired t-test | (n= 6 mTRPV1red transfected animals) | p=0.0189 | | |
| Figure 5K | Paired t-tests | (n= 6 mTRPV1red transfected animals) | pDots=0.0150<br>pLoom=0.4386 | | |
| Figure 5L | Two-Way ANOVA (Repeated-Measures) | (n=6 mTRPV1red transfected animals) | $F_{Interaction}$=6.427 p=0.0011 | Holm-Sidak | Baseline vs Capsaicin:<br>LPR=0-0.5: p=0.8891<br>LPR=0.5-1: p=0.0006<br>LPR=1-2: p=0.0008<br>LPR=2-4: p=0.8891<br>LPR=4+: p=0.9142 |
| Figure 5M | Paired t-test | (n=6 mTRPV1red transfected animals) | p=0.0127 | | |

### Table S6

| Figure | Test Type | Sample Size | Statistics | Multiple Comparisons | Statistics |
|---|---|---|---|---|---|
| Figure 6B | One-Way ANOVA (Repeated-Measures) | (n1=10 animals)<br>(n2=10 animals)<br>(n3=9 animals) | F=11.08<br>p=0.0003 | Holm-Sidak | (1) vs (3): p=0.0003<br>(2) vs (3): p=0.0013 |
| Figure 6C | One-Way ANOVA (Repeated-Measures) | (n1=10 animals)<br>(n2=10 animals)<br>(n3=9 animals) | F=4.846<br>p=0.0163 | Holm-Sidak | (1) vs (3): p=0.0208<br>(2) vs (3): p=0.0144 |
